# The HOG MAPK - Transcription Factor CsAtf1 - *CsErg5B* Regulatory Module Mediates Conidial Germination and Fludioxonil Sensitivity in *Colletotrichum siamense*

**DOI:** 10.64898/2026.05.18.725934

**Authors:** Yuqing Lin, Kuaikuai Wang, Xiaoling Guan, Miao Song, Zhenyu Han, Wenbo Liu, Wei Wu, Yu Zhang, Weiguo Miao, Chunhua Lin

## Abstract

*Colletotrichum siamense* is a predominant causal agent of anthracnose in rubber tree and numerous economically important crops, causing severe yield losses worldwide. Conidial germination represents a critical early step for successful infection, while the high-osmolarity glycerol (HOG) MAPK pathway and ergosterol biosynthesis individually govern fungal development, stress adaptation and fungicide responses. However, the molecular crosstalk between these two modules remains largely elusive in phytopathogenic fungi. Here, we identified *CsErg5B*, a sterol C-22 desaturase homolog, as a direct target of the HOG- regulated transcription factor CsAtf1 in *C. siamense*. *CsErg5B* was indispensable for ergosterol biosynthesis, conidial germination, appressorium formation, and full virulence. The Δ*CsErg5B* mutant showed increased conidiation but severely impaired germination, and exhibited elevated resistance to fludioxonil while hypersensitivity to azole fungicides. Epistasis analysis using the Δ*CsErg5B*/Δ*CsCyp51G1* double mutant - where *CsCyp51G1* serves as another downstream target of CsAtf1 - revealed that *CsErg5B* functions as the predominant downstream effector of CsAtf1 in modulating conidial development and fludioxonil sensitivity. Furthermore, overexpression of *CsErg5B* significantly rescued the defects in conidial germination and fludioxonil sensitivity in both Δ*CsAtf1* and Δ*CsPbs2* mutants. Taken together, our findings uncover a HOG MAPK - CsAtf1 - *CsErg5B* regulatory axis that connects HOG MAPK signaling to ergosterol homeostasis, thereby governing conidial germination and fungicide sensitivity in *C. siamense*. This study provides novel insights into the regulatory network underlying fungal development and fungicide response, and offers promising molecular targets for the integrated management of plant anthracnose.

## 1. Introduction

*Colletotrichum* species are recognized as one of the most economically important phytopathogenic fungi globally, causing destructive anthracnose in diverse hosts, including fruits, tea, rubber trees, and major food and cash crops (Cao et al., 2019; Dofuor et al., 2023; Jeyaraj et al., 2023). Their high epidemic adaptability is largely attributed to abundant conidiation. As major infectious propagules, conidia survive harsh environmental conditions and initiate infection via rapid germination upon host recognition, thereby facilitating polycyclic disease outbreaks (Sephton-Clark and Voelz, 2018; Nordzieke et al., 2019). Conidial germination serves as an essential early prerequisite for host colonization and subsequent disease development (Seong et al., 2008; Jia et al., 2024).

Ergosterol, a predominant sterol in fungal plasma membranes, is essential for maintaining membrane integrity, fluidity, and the activity of membrane-associated proteins, thereby regulating signal transduction and other vital biological processes (Lees et al., 1999; Rodrigues, 2018; Choy et al., 2023). Its biosynthesis is a highly conserved and complex pathway involving approximately 20 enzymes, including well-characterized ones such as Erg9, Erg11 (Cyp51), Erg5, and Erg4 (Lees et al., 1995; Jordá and Puig, 2020). Accumulating evidence indicates that this pathway is pivotal for regulating fungal development and pathogenicity. For instance, in *Magnaporthe oryzae*, disruption of *MoCYP51A* or *MoErg4* impairs conidiation, appressorium formation, and virulence (Yan et al., 2011; Guo et al., 2023). Similarly, deletion of *FgERG3*, *FgERG4*, or *FgERG5* in *Fusarium graminearum* significantly reduces hyphal growth, conidiation, and pathogenicity (Liu et al., 2013; Yun et al., 2014). In *Verticillium dahliae*, *VdERG2* is crucial for ergosterol biosynthesis, vegetative differentiation, and virulence (Lv et al., 2023). Among these enzymes, Erg5, a sterol C-22 desaturase, is essential for normal conidiation and conidial germination in *Aspergillus fumigatus* and *F. oxysporum* (Deng et al., 2015; Long and Zhong, 2022). Despite these findings, the precise molecular mechanisms by which the ergosterol biosynthesis pathway, particularly *Erg5*, governs conidial development and pathogenicity remain largely elusive.

Fungi rely on conserved mitogen-activated protein kinase (MAPK) cascades to cope with complex environments (Saito, 2010; Martínez-Soto and Ruiz-Herrera, 2017). The high-osmolarity glycerol (HOG) MAPK pathway functions as a central stress signaling module via the canonical MAPKKK (Ssk2, Ssk22, Ste11) - MAPKK (Pbs2) - MAPK (Hog1) phosphorylation cascade (de Nadal and Posas, 2022; Tatebayashi and Saito, 2023). This pathway not only mediates stress adaptation, but also serves as a vital and conserved regulator of conidial germination (Jiang et al., 2018; Yaakoub et al., 2022). In the plant pathogen *Alternaria alternata*, *hogA* deletion markedly delays conidial germination (Igbalajobi et al., 2020). Likewise, disruption of HOG signaling or its downstream regulators impairs conidial germination in entomopathogenic *Metarhizium acridum*, human pathogenic *A. fumigatus* and nematode-trapping fungi *Arthrobotrys oligospora* (Jin et al., 2012; Hérivaux et al., 2022; Liu et al., 2025). Collectively, these findings demonstrate that the HOG pathway acts as a core conserved regulator of conidial germination in diverse fungi. Accumulating evidence indicates functional crosstalk between HOG signaling and ergosterol biosynthesis. In yeast and pathogenic fungi such as *Cryptococcus neoformans* and *Candida albicans*, the HOG pathway modulates ergosterol synthetic gene expression to maintain lipid and membrane homeostasis, while ergosterol synthesis inhibition conversely activates the HOG pathway (Ko et al., 2009; Montañés et al., 2011; Tanigawa et al., 2012; Herrero-de-Dios et al., 2020). Both HOG signaling and ergosterol metabolism are essential for conidial germination, yet their coordinated regulatory mechanism during this process remains poorly characterized.

Fludioxonil, a phenylpyrrole fungicide, is widely used for its broad-spectrum efficacy against various fungal pathogens (Wang et al., 2022). Its antifungal activity is primarily attributed to the hyperactivation of the HOG MAPK pathway, which triggers aberrant signal transduction and cellular dysregulation (Proft et al., 2001; Jacob et al., 2014). Downstream transcription factors, particularly the ATF/CREB family, mediate specific cellular responses to the HOG MAPK cascade (Lara-Rojas et al., 2011; Hagiwara et al., 2014). As a core member of this family, ATF1 functions downstream of Hog1 to govern oxidative stress tolerance and full pathogenicity (Li et al., 2023). In our previous work, we demonstrated that the CsAtf1 ortholog in *C. siamense* regulates conidial morphogenesis, cell wall integrity, fludioxonil sensitivity and pathogenicity (Song et al., 2022). We further revealed that CsAtf1 negatively regulates the cytochrome P450 gene *CsCyp51G1*, thereby enhancing fungal sensitivity to fludioxonil (Guan et al., 2022). However, the link between the ergosterol biosynthesis pathway and fungicide sensitivity remains unclear, and the molecular mechanisms by which the HOG pathway regulates these processes through downstream transcription factors, including whether it directly controls the ergosterol biosynthesis pathway, remain poorly understood.

In this study, we identified *CsErg5B*, a conserved ortholog encoding sterol C-22 desaturase, as a direct target gene of the transcription factor CsAtf1 in *C. siamense*. We demonstrated that *CsErg5B* is essential for ergosterol biosynthesis and plays a pivotal role in conidial germination, appressorium formation, fludioxonil sensitivity, and full virulence. Using a Δ*CsErg5B*/Δ*CsCyp51G1* double deletion mutant, we provided genetic evidence that *CsErg5B* acts as the dominant effector of CsAtf1 for these traits. Furthermore, CsAtf1 regulated these functions by modulating *CsErg5B* expression, and overexpression of *CsErg5B* in both Δ*CsAtf1* and the upstream Δ*CsPbs2* mutants rescued the defects in conidial germination and fludioxonil sensitivity, positioning *CsErg5B* as a key downstream effector of HOG MAPK pathway. This work uncovers a novel HOG MAPK - CsAtf1 - *CsErg5B* regulatory axis linking HOG MAPK and ergosterol biosynthesis in regulating conidial germination and fludioxonil sensitivity, shedding new light on the molecular mechanisms governing fungicide sensitivity and infection-related morphogenesis in *C. siamense*.

## 2 Results

### 2.1 *CsErg5B* is a target gene positively regulated by the transcription factor CsAtf1

Previous studies have identified CsAtf1 as a key downstream transcription factor of the HOG MAPK pathway (Song et al., 2022). Chromatin immunoprecipitation sequencing (ChIP-Seq) and high-throughput RNA-sequencing (RNA-Seq) analyses screened potential target genes directly modulated by CsAtf1, among which *CsErg5B* was significantly downregulated in the Δ*CsAtf1* mutant (Guan et al., 2022). The full-length coding sequence of *CsErg5B* was amplified using the primer pair *CsErg5B*-F/R and deposited in GenBank (Accession No. OP677558). Sequence analysis showed that the 1770-bp genomic region contains three introns, and the 1602-bp cDNA encodes a 533-amino-acid protein harboring a conserved cytochrome P450 domain (Pfam; E-value: 3.6e-48) (Fig S1).

To verify the direct regulatory relationship between CsAtf1 and *CsErg5B*, yeast one-hybrid (Y1H) assays were performed. The full-length cDNA of *CsAtf1* was cloned into the GAL4 activation domain vector (pGADT7-*CsAtf1*), and the 1500-bp promoter region of *CsErg5B* was inserted into the pHIS2 reporter vector. Co-transformation of these constructs into yeast strain Y187 enabled growth on SD/-Trp/-Leu/-His medium supplemented with 80 mM 3-AT (to suppress auto-activation), whereas control cells co-transformed with empty vectors failed to grow under the same conditions (Fig 1A). This direct interaction was further validated via EMSA. The recombinant CsAtf1 protein bound to the *CsErg5B* promoter and formed stable DNA-protein complexes, leading to a distinct mobility shift. The specificity of this binding was confirmed by efficient competition with unlabeled competitor probes (Fig 1B).

**Fig 1.**
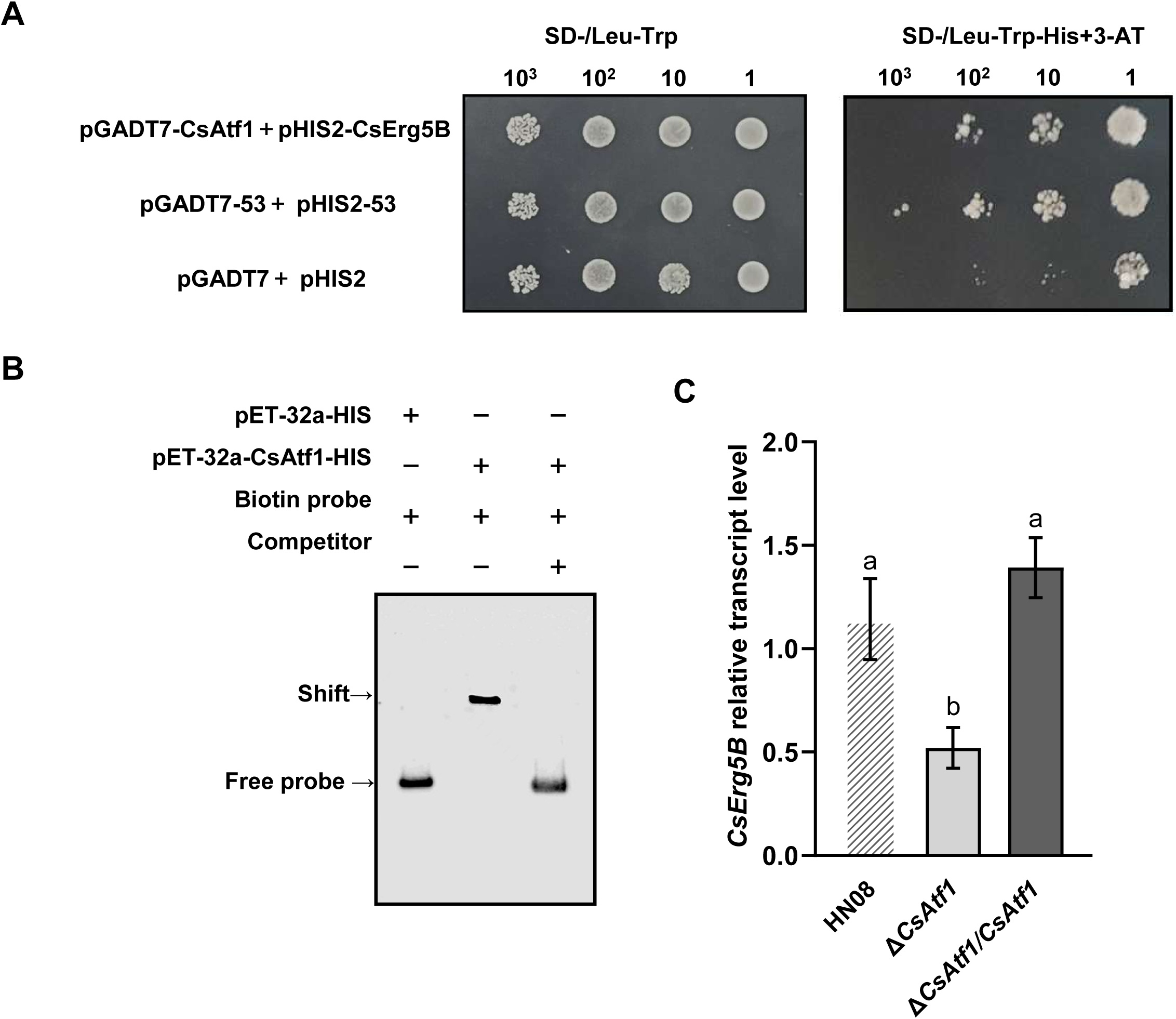
Binding and expression analysis of CsAtf1 with *CsErg5B*. **(A)** Yeast one-hybrid assay for interaction between CsAtf1 and the *CsErg5B* promoter. 10-fold serial dilutions (10³ to 1) of transformed yeast cells were spotted on SD/−Leu/−Trp and SD/−Leu/−Trp/−His supplemented with 80 mM 3-AT selective media. **(B)** EMSA confirmed that CsAtf1 binds to the *CsErg5B* promoter in vitro. “+”/“−” denote presence/absence of components. Shifted complex and free probe are labeled. **(C)** Relative transcript levels of *CsErg5B* in wild-type HN08, Δ*CsAtf1* mutant, and the complemented strain Δ*CsAtf1*/*CsAtf1*. Data are mean ± SD. Different letters indicate significant differences at *p* < 0.05 according to one-way ANOVA and Duncan’s test.

To further confirm the transcriptional regulation of *CsErg5B* by CsAtf1, we examined *CsErg5B* transcript levels in wild-type strain HN08, Δ*CsAtf1* mutant and complemented strain using qRT-PCR. The expression of *CsErg5B* was markedly reduced by nearly 50% in the Δ*CsAtf1* mutant compared with the wild type (Fig 1C). Collectively, these results demonstrate that CsAtf1 directly binds to the *CsErg5B* promoter and positively regulates its expression.

### 2.2 *CsErg5B* is involved in conidial germination, appressorium formation and ergosterol biosynthesis

To investigate the biological function of *CsErg5B*, we generated a gene deletion mutant (Δ*CsErg5B*) and overexpression strains (Δ*CsErg5B*/*CsErg5B*-OE), which were verified by PCR, sequencing, Southern blot analysis, and qRT-PCR (Fig S2). Phenotypic characterization demonstrated that *CsErg5B* differentially regulates vegetative growth and asexual reproduction. Colony diameters exhibited modest alterations: Δ*CsErg5B* increased by 3.04%, whereas Δ*CsErg5B*/*CsErg5B*-OE decreased by 2.30% relative to the wild-type (Fig 2A, 2B). In contrast, conidiation was dramatically affected: Δ*CsErg5B* produced approximately 2.65-fold more conidia (10.3 × 10⁵ conidia/mL) than the wild-type (2.83 × 10⁵ conidia/mL), while Δ*CsErg5B*/*CsErg5B*-OE showed a slight reduction (9.40%) (Fig 2C).

**Fig 2.**
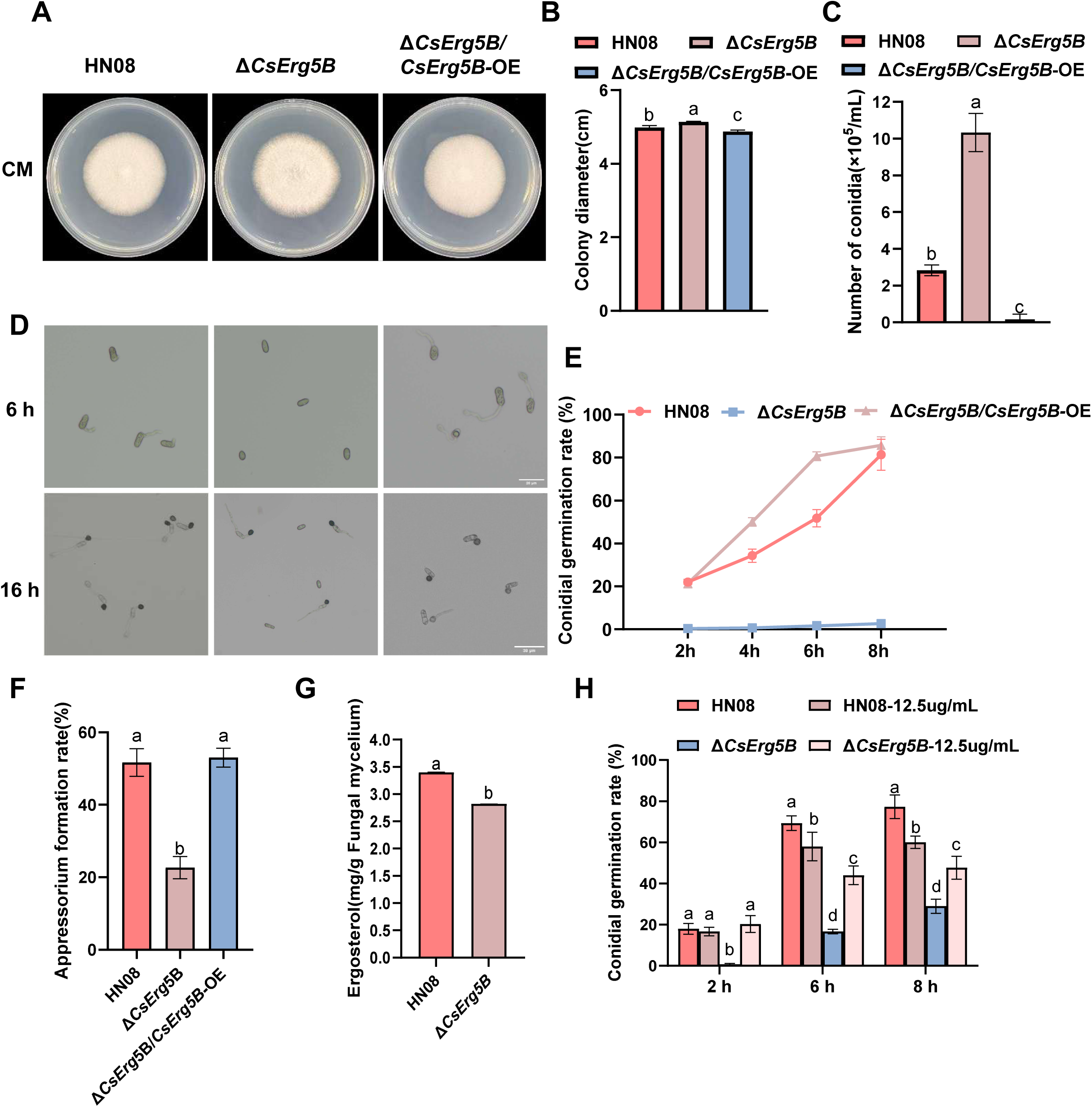
Roles of *CsErg5B* in conidial development and ergosterol synthesis. **(A, B)** Colony diameter of each strain after cultivation. **(C)** Quantitative detection of conidiation among different strains. **(D, E)** Conidial germination rates at 2, 4, 6 and 8 hpi. **(D, F)** Appressorium formation rates at 16 hpi. **(G)** Determination of intracellular ergosterol content by HPLC. **(H)** Effects of exogenous ergosterol on conidial germination rate were determined at 2, 6, and 8 hpi. Wild-type HN08 and Δ*CsErg5B* conidia were cultured with or without 12.5 μg/mL exogenous ergosterol. Data are mean ± SD. Different letters indicate significant differences at *p* < 0.05 according to one-way ANOVA and Duncan’s test.

Strikingly, despite its hyper-conidiation phenotype, Δ*CsErg5B* exhibited severe defects in conidial germination, with significantly lower germination rates at all examined time points 2-8 hours post-inoculation (hpi) compared to both the wild-type and Δ*CsErg5B*/*CsErg5B*-OE (Fig 2D, 2E). Similarly, appressorium formation was markedly impaired in Δ*CsErg5B* (22.67% at 16 hpi) compared with the wild-type (51.67%) and Δ*CsErg5B*/*CsErg5B*-OE (53.00 %) (Fig 2D, 2F). Consistent with its predicted role in sterol biosynthesis, Δ*CsErg5B* showed a 17.06% reduction in ergosterol content (2.82 mg/g) relative to the wild-type (3.40 mg/g) as determined by HPLC (Fig 2G). To determine whether the conidial germination defect in Δ*CsErg5B* is causally linked to ergosterol deficiency, we examined the effect of exogenous ergosterol supplementation on conidial germination in the wild-type (HN08) and Δ*CsErg5B* strains. Notably, exogenous supplementation with 12.5 μg/mL ergosterol partially rescued the conidial germination defect of the Δ*CsErg5B* mutant, while slightly suppressing germination in the wild-type strain (Fig 2H). Taken together, these findings indicate that *CsErg5B* governs ergosterol biosynthesis, conidial germination competence and appressorium formation, and regulates conidial germination via the ergosterol signaling pathway.

### 2.3 *CsErg5B* is indispensable for full virulence

Given the impaired infection-related morphogenesis in Δ*CsErg5B*, we evaluated its virulence on rubber tree leaves. Conidial suspensions (1 × 10^6^ conidia/mL) of the wild-type HN08, Δ*CsErg5B*, and Δ*CsErg5B*/*CsErg5B*-OE were inoculated onto detached leaves with or without wounding. After 5 days post-inoculation, Δ*CsErg5B* caused significantly smaller lesions compared to the wild-type under both conditions. On wounded leaves, the lesion areas as 0.44 ± 0.22 cm² for Δ*CsErg5B*, compared to 0.81 ± 0.47 cm² for the wild-type and 1.18 ± 0.54 cm² for Δ*CsErg5B*/*CsErg5B*-OE (Fig 3A, 3B). On unwounded leaves, the lesion areas were 0.31 ± 0.15 cm², 0.69 ± 0.30 cm², and 0.83 ± 0.25 cm² for Δ*CsErg5B*, the wild-type, and Δ*CsErg5B*/*CsErg5B*-OE, respectively (Fig 3C, 3D). These results indicate that *CsErg5B* is required for full virulence in *C. siamense*.

**Fig 3.**
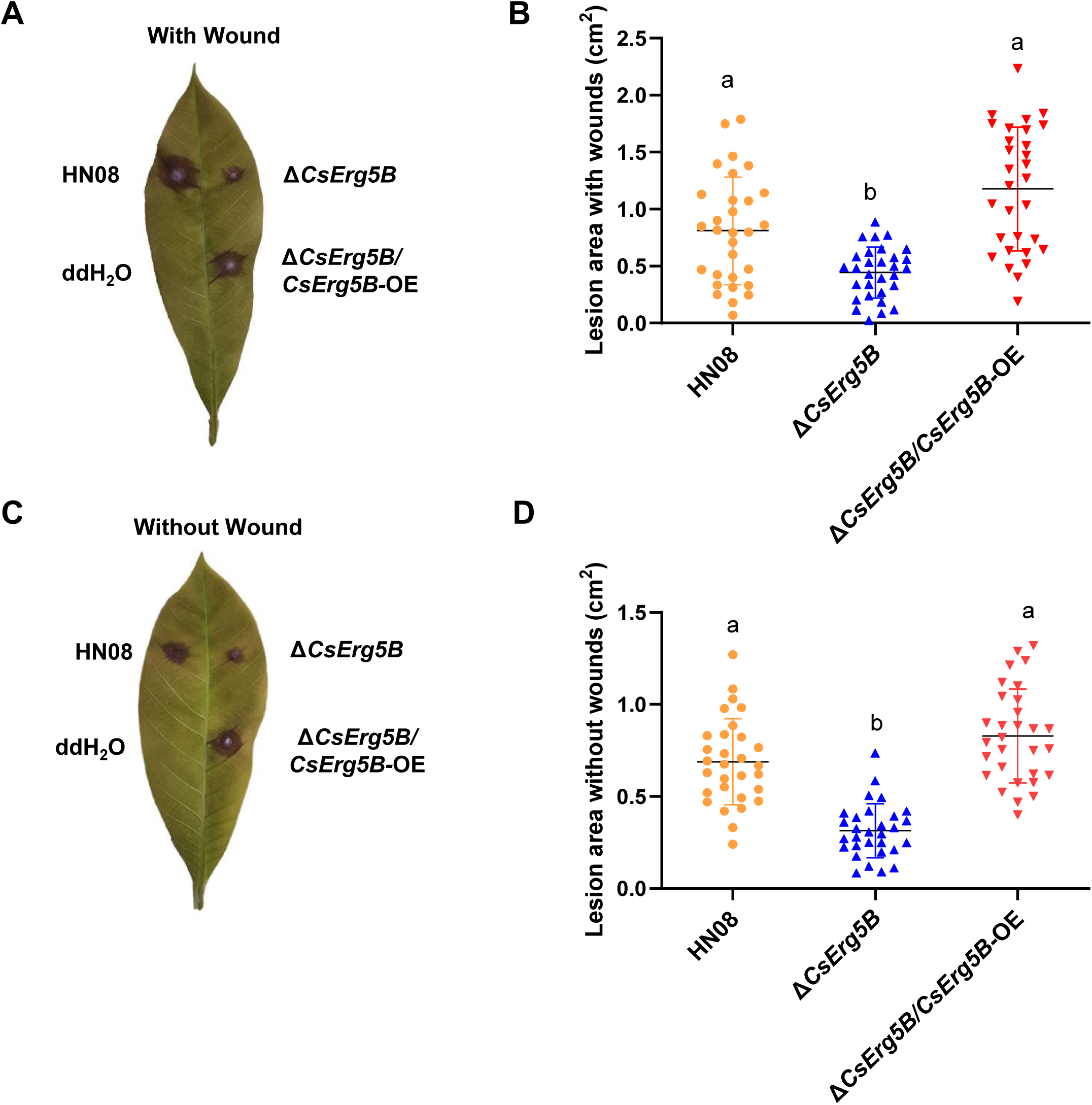
*CsErg5B* deletion attenuates virulence in *C. siamense*. **(A, B)** Disease lesions and statistical analysis of lesion areas on wounded leaves at 5 days post-inoculation. **(C, D)** Pathogenic phenotypes and lesion size quantification on unwounded leaves. Different letters indicate significant differences at *p* < 0.05 according to one-way ANOVA and Duncan’s test.

### 2.4 Deletion of *CsErg5B* affects stress responses and sensitivity to fungicides

To assess the role of *CsErg5B* in stress adaptation, we compared the stress responses of the wild-type strain HN08, Δ*CsErg5B*, and Δ*CsErg5B*/*CsErg5B*-OE under various conditions. No significant differences were observed among strains under salt stress (0.6 M NaCl, 0.7 M KCl) or cell wall stress (50 μg/mL Congo Red), indicating that *CsErg5B* does not affect cell wall integrity or salt stress responses (Fig S3A, S3B). However, Δ*CsErg5B* exhibited reduced sensitivity to osmotic stress induced by 1 M sorbitol (inhibition rates: 9.83% for HN08, 4.95% for Δ*CsErg5B*, 8.63% for Δ*CsErg5B*/*CsErg5B*-OE) and 1 M glucose (17.92%, 10.51%, and 16.26%, respectively) (Fig S3A, S3B). Notably, Δ*CsErg5B* showed enhanced sensitivity to oxidative stress: under 2.5 mM H₂O₂, the inhibition rates were 24.30% for HN08, 50.79% for Δ*CsErg5B*, and 24.94% for Δ*CsErg5B/CsErg5B*-OE (Fig S3A, S3B). These results reveal distinct roles for *CsErg5B* in osmotic and oxidative stress adaptation.

We next examined fungicide sensitivity. With respect to azole fungicides, Δ*CsErg5B* exhibited significantly enhanced sensitivity to tebuconazole (0.25-2 μg/mL) and difenoconazole (0.01-1 μg/mL) compared to the wild-type and Δ*CsErg5B*/*CsErg5B*-OE (Fig 4A-4D). Conversely, under fludioxonil stress (0.01-5 μg/mL), Δ*CsErg5B* showed significantly stronger resistance than the wild-type, while Δ*CsErg5B*/*CsErg5B*-OE displayed enhanced sensitivity (Fig 4E, 4F). These results demonstrate that *CsErg5B* positively regulates azole resistance and fludioxonil sensitivity in *C. siamense*, suggesting that *CsErg5B* plays opposing regulatory roles in responses to different classes of fungicides.

**Fig 4.**
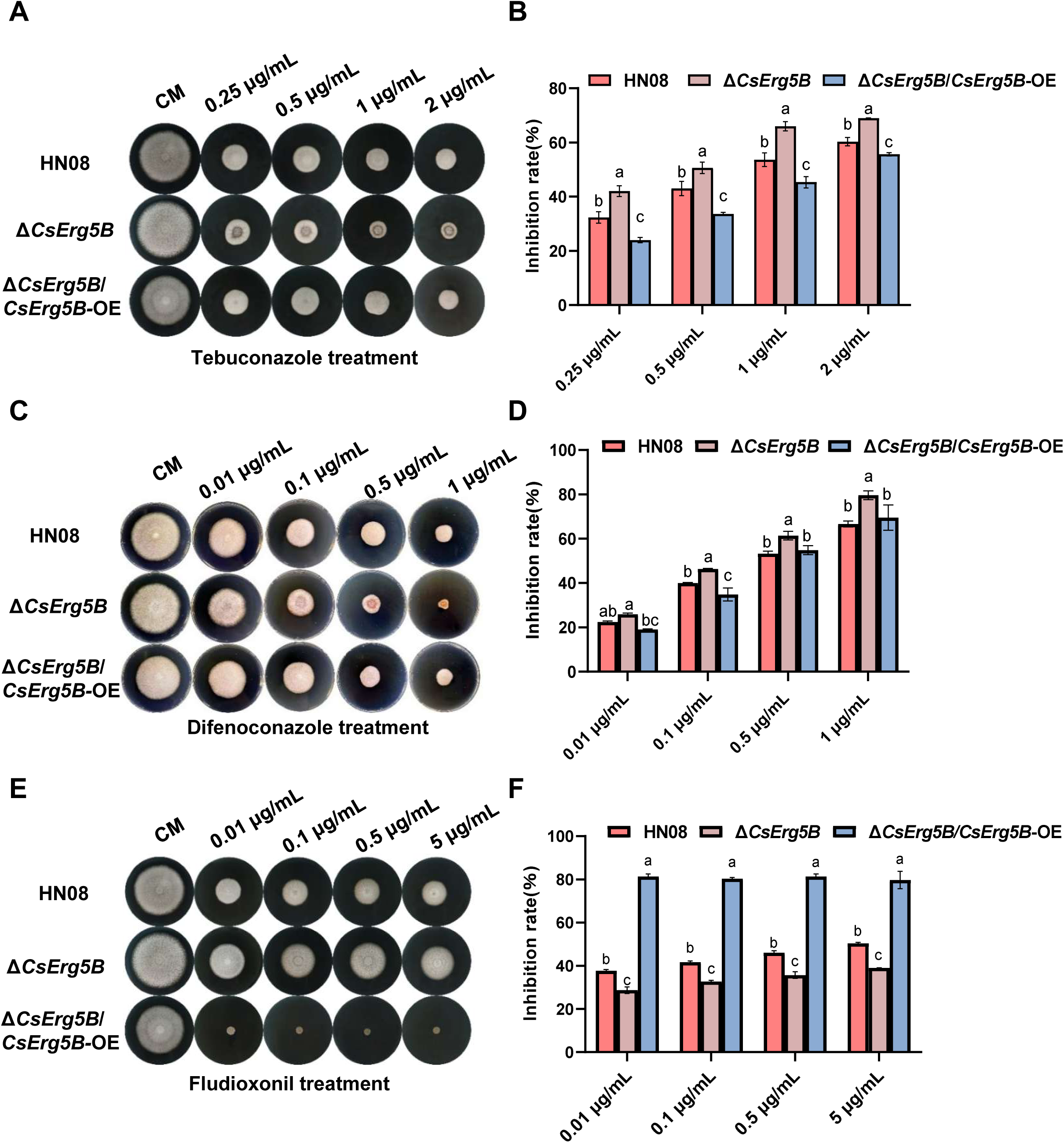
Effects of *CsErg5B* deletion on fungicide sensitivity. **(A–D)** Sensitivity determination to azole fungicides tebuconazole and difenoconazole. **(E, F)** Mycelial growth inhibition under fludioxonil exposure. Data are mean ± SD. Different letters indicate significant differences at *p* < 0.05.

### 2.5 *CsErg5B*, rather than *CsCyp51G1,* acts as the dominant downstream effector of CsAtf1

As reported in our previous study, *CsCyp51G1*, a target gene negatively regulated by the transcription factor CsAtf1, contributed to increased sensitivity to fludioxonil upon its deletion (Guan et al., 2022). In contrast, deletion of *CsErg5B* in the present study resulted in decreased sensitivity to fludioxonil. To determine which gene plays a predominant role in these phenotypes, a double deletion mutant Δ*CsErg5B*/Δ*CsCyp51G1* was generated (Fig S4). The results showed that conidial germination rates of the Δ*CsErg5B*/Δ*CsCyp51G1* double mutant at 2-8 hpi were lower than those of the wild type, the single deletion mutants Δ*CsCyp51G1* and Δ*CsErg5B*, and were more similar to the phenotype of Δ*CsErg5B* (Fig 5A, 5B). In addition, the appressorium formation rate of the double mutant at 16 hpi was 0%, which was significantly lower than that of the wild type (54.00%), Δ*CsErg5B* (15.33%), and Δ*CsCyp51G1* (64.67%) (Fig 5A, 5C).

**Fig 5.**
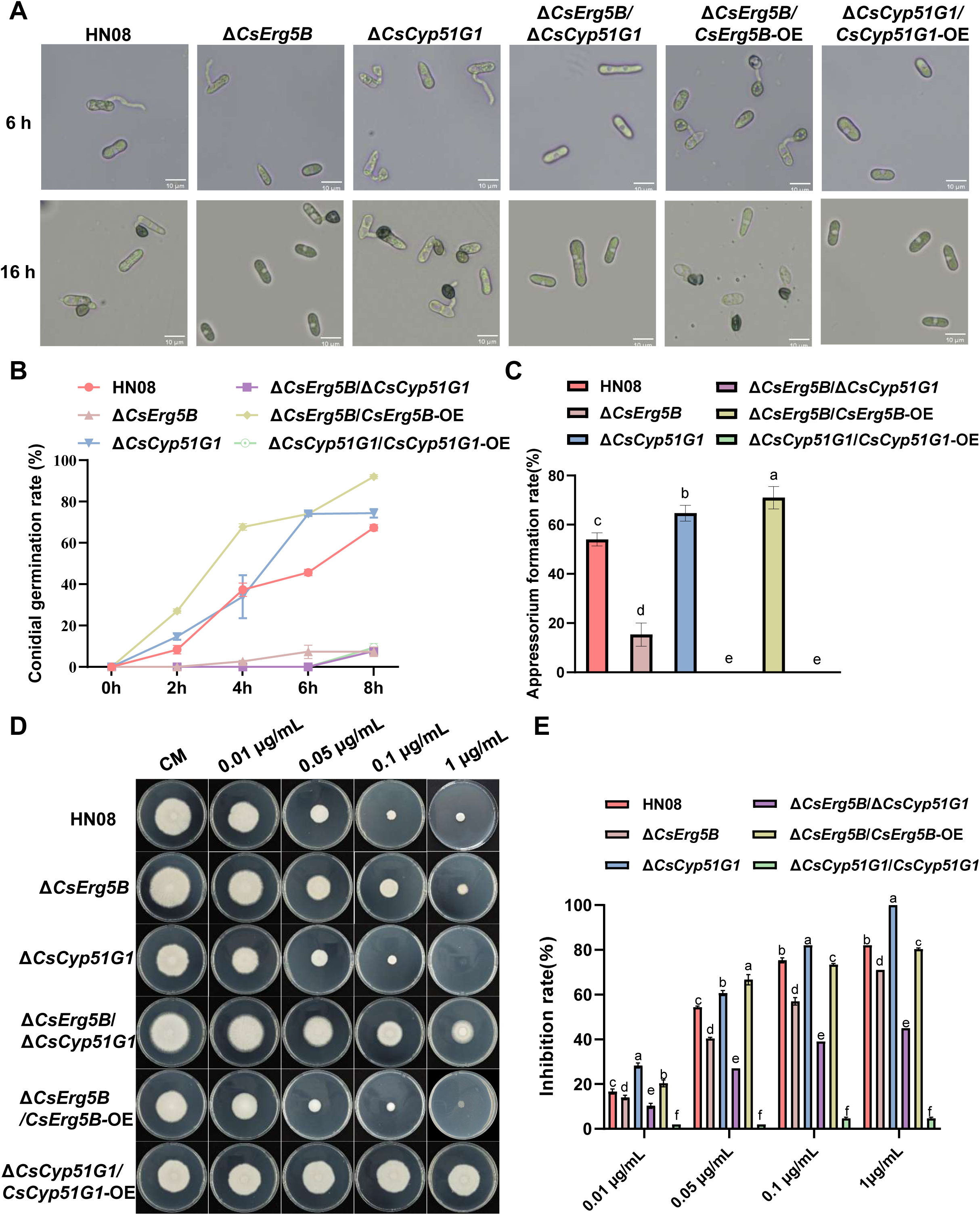
Conidial Germination, appressorium formation and fludioxonil sensitivity in Δ*CsErg5B*/Δ*CsCyp51G1* double mutant. **(A-C)** Conidial germination rates at 2-8 hpi and appressorium formation rates at 16 hpi were statistically analyzed the wild-type HN08, single mutants (Δ*CsErg5B*, Δ*CsCyp51G1*), corresponding overexpression strains, and double mutant Δ*CsErg5B*/Δ*CsCyp51G1* were examined. **(D-E)** Fludioxonil tolerance of different strains were evaluated under 0.01–1.00 μg/mL fludioxonil treatments.Data are mean ± SD. Different letters indicate significant differences at *p* < 0.05.

In terms of fludioxonil sensitivity, the Δ*CsErg5B*/Δ*CsCyp51G1* double mutant exhibited enhanced resistance compared with the wild type when treated with fludioxonil at concentrations ranging from 0.01 to 1.00 μg/mL. This resistance phenotype was consistent with that of the Δ*CsErg5B* single mutant but completely opposite to that of the Δ*CsCyp51G1* single mutant, which showed increased sensitivity to the same concentration gradient of fludioxonil (Fig 5D, 5E). Collectively, these results demonstrate that among the two CsAtf1-regulated target genes *CsCyp51G1* and *CsErg5B*, *CsErg5B* acts as the dominant effector gene governing conidial germination, appressorium formation, and fludioxonil sensitivity.

### 2.6 Overexpression of *CsErg5B* rescues conidial germination and fludioxonil sensitivity defects in the Δ*CsAtf1* mutant

To explore the regulatory hierarchy of the HOG MAPK pathway, we analyzed the expression of *CsPbs2*, *CsHog1*, *CsAtf1* and *CsErg5B* in *C. siamense* under 2×10^−3^ μg/mL anisomycin (a HOG MAPK activator), 20 μg/mL SB203580 (a p38 MAPK inhibitor), and 50 μg/mL fludioxonil (a phenylpyrrole fungicide) (Fig S5). The four genes were significantly activated by the HOG pathway activator anisomycin and fludioxonil, but repressed by the pathway inhibitor SB203580, indicating that these genes are co-regulated under HOG pathway modulation, consistent with a HOG MAPK -CsAtf1- *CsErg5B* regulatory cascade in *C. siamense*.

Given that the Δ*CsAtf1* mutant exhibits phenotypic defects, including impaired conidial germination, appressorium formation, and altered fungicide sensitivity (Song et al., 2022), and these defects resemble those of the Δ*CsErg5B* mutant, we hypothesized that these phenotypes may be mediated through CsAtf1 dependent regulation of *CsErg5B*. To test this hypothesis, we constructed a *CsErg5B* overexpression vector (pBAR-*gpdA*-*CsErg5B*) and introduced it into the Δ*CsAtf1* mutant. The resulting transformants (designated Δ*CsAtf1*/*CsErg5B*-OE) were verified by PCR and qRT-PCR (Fig S6).

Phenotypic analysis revealed that overexpression of *CsErg5B* partially rescued the defects of the Δ*CsAtf1* mutant. At 6 hours post-inoculation, conidial germination rates were 54.00% in wild type (HN08), 31.67% in Δ*CsAtf1*, and 60.00% in Δ*CsAtf1*/*CsErg5B*-OE (Fig 6A, 6B), indicating that *CsErg5B* overexpression restored the germination defect in the Δ*CsAtf1* mutant. However, appressorium formation remained impaired in Δ*CsAtf1*/*CsErg5B*-OE (Fig 6C, 6D), and virulence assays on rubber tree leaves confirmed that *CsErg5B* overexpression failed to restore its pathogenicity, suggesting that additional CsAtf1-dependent factors are required for appressorium formation and virulence (Fig S7).We further assessed fungicide sensitivity by measuring mycelial growth on medium containing fludioxonil (0.01-10 μg/mL). Consistent with previous reports, the Δ*CsAtf1* mutant exhibited reduced sensitivity to fludioxonil compared to the wild-type. Strikingly, *CsErg5B* overexpression in the Δ*CsAtf1* mutant restored fludioxonil sensitivity to levels comparable to the wild-type (Fig 6E, 6F). Taken together, these results demonstrate that *CsErg5B* acts downstream of CsAtf1 to regulate conidial germination and fludioxonil sensitivity.

**Fig 6.**
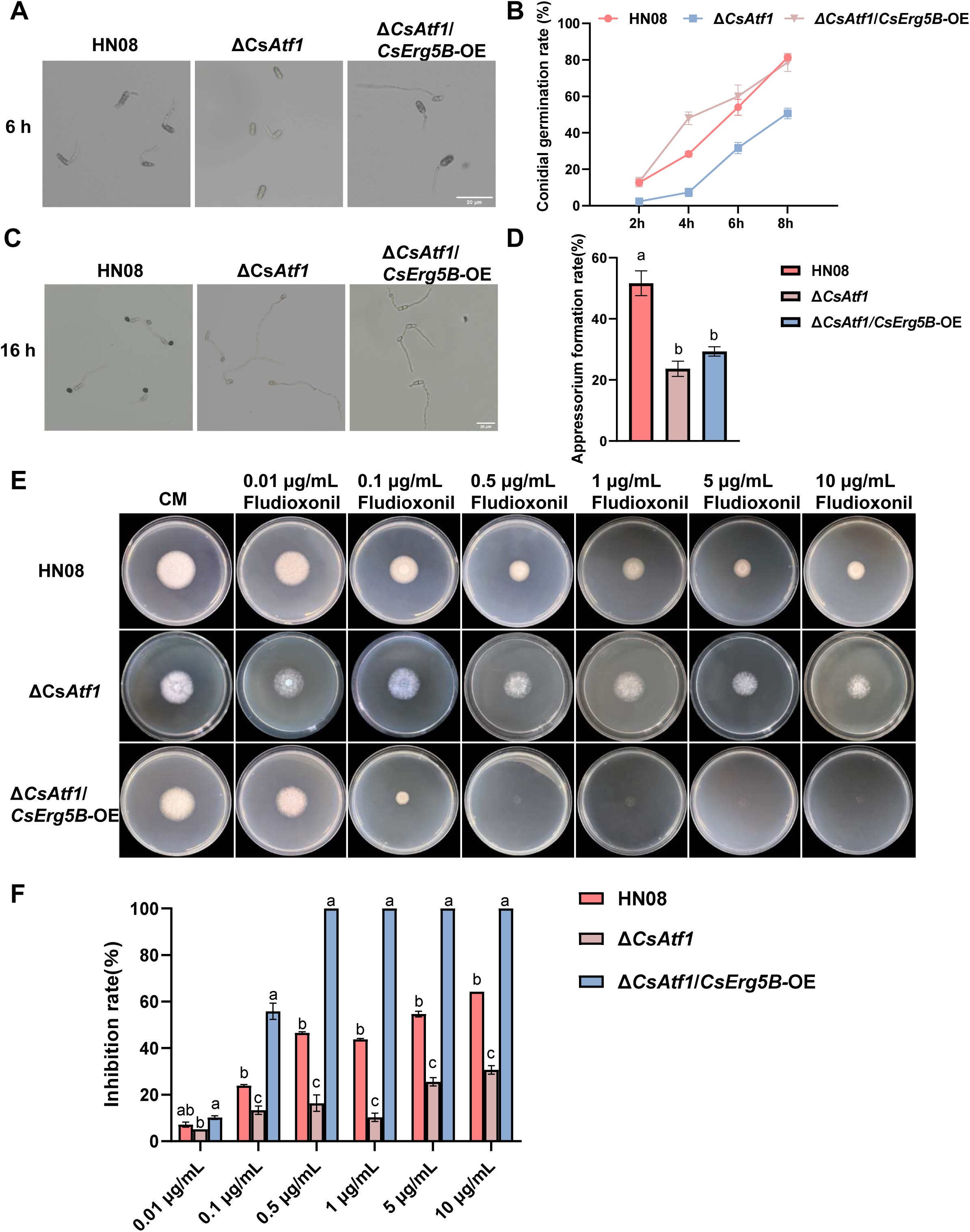
Phenotypic characteristics of the wild-type HN08, Δ*CsAtf1* mutant and Δ*CsAtf1*/*CsErg5B*-OE strain were determined. **(A–D)** Conidial germination at 2, 4, 6 and 8 hpi, as well as appressorium formation at 16 hpi, were examined across the three strains. **(E–F)** Mycelial growth and fludioxonil sensitivity of tested strains were evaluated under 0.01–10.00 μg/mL fludioxonil treatments. All data are presented as mean ± SD. Different letters indicate significant differences at *p* < 0.05.

### 2.7 Overexpression of *CsErg5B* rescues conidial germination and fludioxonil sensitivity defects in the Δ*CsPbs2* mutant

Previously, we generated a gene deletion mutant of *CsPbs2* (encodes a key MAPKK component of the HOG pathway) that. significantly impairs conidial germination and fludioxonil sensitivity in *C. siamense* (Lin et al., 2018). However, the molecular basis for these defects remained unclear. To determine whether they arise from dysregulated control of *CsErg5B*, we constructed a *CsErg5B* overexpression vector and transformed it into the Δ*CsPbs2* mutant. The resulting positive transformants (designated Δ*CsPbs2*/*CsErg5B*-OE) were verified by PCR and qRT- PCR analyses (Fig S6C, S6D).

Conidial germination was assessed at 2-8 hpi in wild-type, Δ*CsPbs2*, and Δ*CsPbs2*/*CsErg5B*-OE strains. Overexpression of *CsErg5B* effectively rescued the conidial germination defect in the Δ*CsPbs2* mutant (Fig 7A, 7B). In contrast, appressorium formation was unaffected by either *CsPbs2* deletion or *CsErg5B* overexpression in the Δ*CsPbs2* background (Fig 7C, 7D), indicating that this developmental process is regulated independently of the Pbs2- mediated *Erg5B* signaling.Fludioxonil sensitivity was evaluated by measuring colony growth on CM medium containing fludioxonil (0.01-10 μg/mL) after 3 days of incubation. Notably, *CsErg5B* overexpression restored fludioxonil sensitivity in the Δ*CsPbs2* mutant to near wild-type levels (Fig 7E, 7F). Collectively, these findings demonstrate that CsPbs2 regulates conidial germination and fludioxonil sensitivity, at least in part, through CsErg5B.

**Fig 7.**
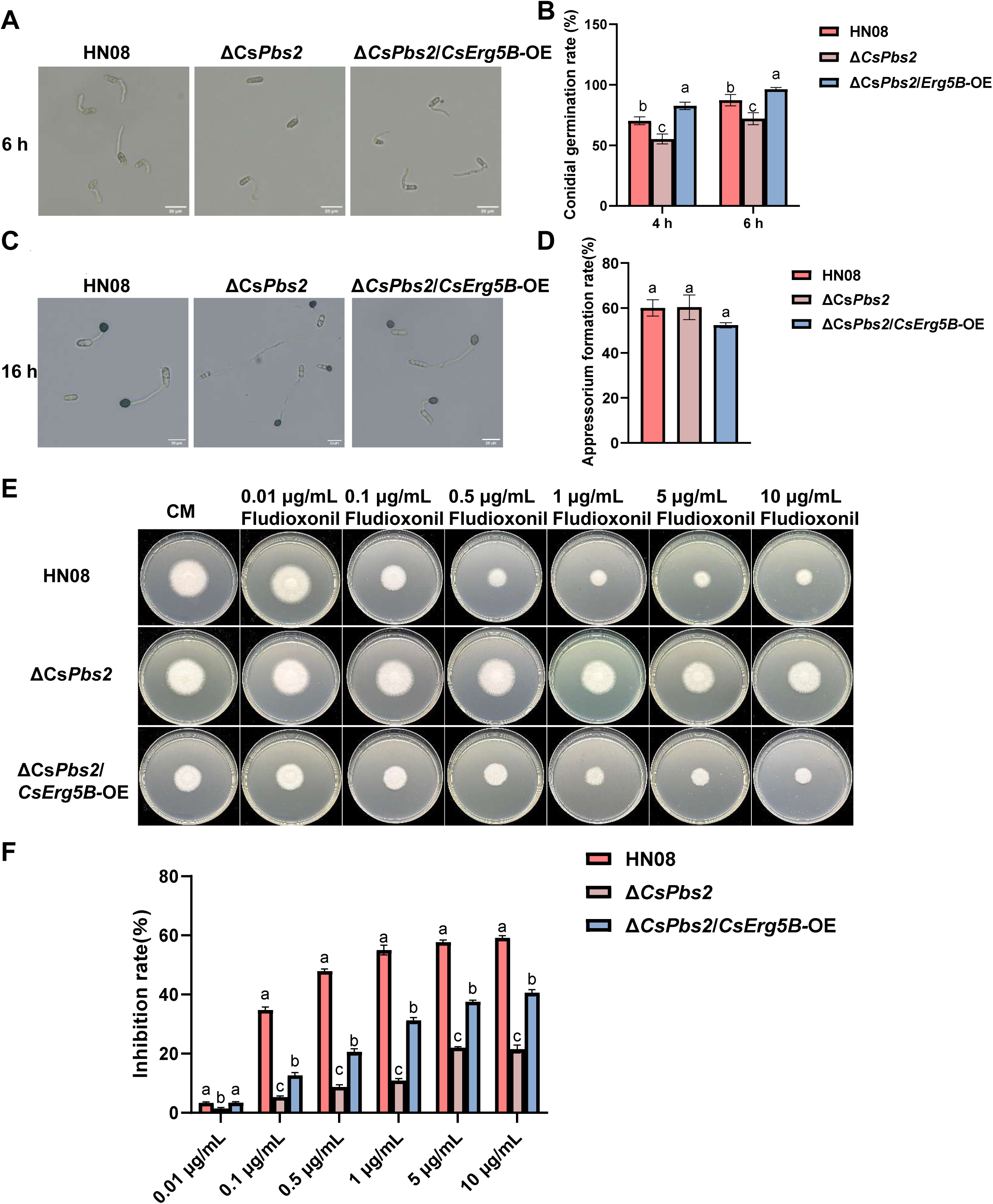
Phenotypic characteristics of the wild-type HN08, Δ*CsPbs2* mutant and Δ*CsPbs2*/*CsErg5B*-OE strain were determined. **(A–D)** Conidial germination at 2, 4, 6 and 8 hpi, as well as appressorium formation at 16 hpi, were examined across the three strains. **(E–F)** Mycelial growth and fludioxonil sensitivity of tested strains were evaluated under 0.01–10.00 μg/mL fludioxonil treatments. All data are presented as mean ± SD. Different letters indicate significant differences at *p* < 0.05.

## 3. Discussion

The HOG MAPK pathway is well known to govern multiple stress responses and development in plant pathogenic fungi, but how it crosstalks with ergosterol biosynthesis to modulate conidial germination and fungicide sensitivity remains poorly understood (Ji et al., 2012; Ma et al., 2022; Zhao et al., 2022; Galello et al., 2024). In this study, we demonstrated that *CsErg5B*, an ortholog encoding sterol C-22 desaturase essential for ergosterol biosynthesis, is directly targeted by the transcription factor CsAtf1 and acts as a key downstream effector of the HOG pathway. Furthermore, we identified a novel HOG MAPK - CsAtf1 - *CsErg5B* regulatory axis that mediates conidial germination andfludioxonil sensitivity in *C. siamense*. This work fills a critical knowledge gap by establishing a mechanistic link between HOG MAPK signaling and ergosterol metabolism in regulating both processes.

Ergosterol is a core component of fungal cell membranes, and its biosynthetic pathway directly determines membrane integrity, fluidity, and membrane protein function. Disruption of key enzymes in this pathway causes growth arrest, conidiation defects, or even lethality, underscoring its essential role in fungal survival (Fang et al., 2023; Guo et al., 2023). For instance, sterol 14α-demethylase in *M. oryzae* modulates conidiation, virulence, and sensitivity to sterol demethylation inhibitors. *V. dahliae VdERG2* and *P. expansum Erg4* are required for ergosterol biosynthesis and normal fungal development (Yan et al., 2011; Han et al., 2023; Lv et al., 2023). As a sterol C-22 desaturase, Erg5 is highly conserved in pathogenic fungi including *A. fumigatus*, *F. graminearum* and *C. albicans*, where it mediates ergosterol biosynthesis, vegetative differentiation, and virulence (Sun et al., 2013; Yun et al., 2014; Long and Zhong, 2022). Consistent with these reports, deletion of *CsErg5B* in *C. siamense* reduced ergosterol levels by 17% and severely impaired conidial germination and appressorium formation. Notably, the Δ*CsErg5B* mutant displayed hyper-conidiation, in contrast to the reduced conidiation caused by *erg5* deletion in other fungi (Yun et al., 2014; Long and Zhong, 2022). Thus, *CsErg5B* differentially regulates asexual development: it negatively controls conidial number but positively maintains conidial quality and germination competence. This trade-off suggests a species-specific regulatory role in *C. siamense*. Exogenous ergosterol supplementation partially restored the conidial germination defect in Δ*CsErg5B* (Fig 2H), confirming that the germination impairment is causally linked to ergosterol deficiency. The partial rescue may reflect limited uptake or incorporation of exogenous sterol into the plasma membrane, or the presence of additional ergosterol- independent germination pathways.

Given the essential role of the ergosterol biosynthetic pathway in fungal growth and membrane homeostasis, this pathway has long been recognized as a major target of antifungal agents, especially azole compounds (Baker et al., 2022; Herrick et al., 2026). As a key enzyme in this pathway, *CsErg5B* exhibited a striking dual role in regulating fungal sensitivity to different classes of fungicides, an intriguing finding of this study. Specifically, deletion of *CsErg5B* conferred resistance to fludioxonil, while increasing susceptibility to azole fungicides including tebuconazole and difenoconazole (Fig 4). This divergent response likely reflects the distinct modes of action of these fungicides. Fludioxonil exerts antifungal activity by hyperactivating the HOG MAPK pathway, leading to cellular dysfunction (Proft et al., 2001; Jacob et al., 2014). The increased fludioxonil resistance in Δ*CsErg5B* may result from altered membrane properties caused by reduced ergosterol levels, which could affect drug uptake or signal transduction. In contrast, azole fungicides directly target ergosterol biosynthesis by inhibiting the CYP51 (Erg11) enzyme (Van Rhijn et al., 2021; Pérez-Cantero et al., 2023; Ishii et al., 2025). The heightened azole sensitivity in Δ*CsErg5B* can be attributed to synthetic lethality, caused by the combined effect of genetic blockage and chemical inhibition of ergosterol biosynthesis, whereas similar phenomena have been documented in other fungi (Baker et al., 2022; Dladla et al., 2024; Saha et al., 2024). Consistent with a link to HOG signaling, overexpression of *CsErg5B* in both Δ*CsAtf1* and Δ*CsPbs2* backgrounds restored fludioxonil sensitivity (Figs 6, 7). This result further confirms that *CsErg5B* is a critical determinant of fungicide response downstream of HOG MAPK signaling.

Importantly, to determine which of the two CsAtf1-regulated ergosterol genes, *CsErg5B* or *CsCyp51G1*, plays the dominant role in the observed phenotypes, we constructed a Δ*CsErg5B*/Δ*CsCyp51G1* double deletion mutant. Phenotypic analysis of this double mutant clearly showed that its conidial germination rate, appressorium formation, and fludioxonil sensitivity were almost identical to those of the Δ*CsErg5B* single mutant, rather than resembling the Δ*CsCyp51G1* single mutant (Fig 5). Specifically, the double mutant exhibited severely impaired germination (similar to Δ*CsErg5B*), a complete loss of appressorium formation (0 % at 16 hpi), and enhanced resistance to fludioxonil (opposite to the hypersensitive phenotype of Δ*CsCyp51G1*). These results unambiguously demonstrate that *CsErg5B* acts as the dominant effector downstream of CsAtf1 for these traits, and that the contribution of *CsCyp51G1* is epistatically masked by *CsErg5B* under the tested conditions. This finding refines our understanding of the CsAtf1 regulatory network: although CsAtf1 controls multiple target genes, *CsErg5B* is the principal mediator of conidial germination and fludioxonil sensitivity.

After clarifying the functions of *CsErg5B* in ergosterol biosynthesis, asexual development, and fungicide responses in *C. siamense*, we focused on its transcriptional regulation. Although ergosterol biosynthetic enzymes have been well studied, the transcriptional regulatory mechanisms of key pathway genes remain poorly understood, with only a few regulators and pathways identified to maintain ergosterol homeostasis. Current understanding of ergosterol gene transcription mainly comes from model and pathogenic fungi. In *Saccharomyces cerevisiae*, zinc-cluster transcription factors Upc2 and Ecm22 activate ergosterol genes under sterol depletion (Vik A and Rine, 2001). Under hyperosmotic stress, the Hog1 MAP kinase regulates *ERG* genes via repressors Mot3 and Rox1, linking stress signaling to sterol metabolism (Martínez-Soto and Ruiz-Herrera, 2017). Consistent with these observations, the HOG pathway is closely linked to ergosterol biosynthesis across diverse fungi. In *C. neoformans*, mutations in HOG pathway components (Hog1, Pbs2, Ssk1) alter ergosterol levels and azole sensitivity (Bahn and Jung, 2013). In *F. graminearum*, tebuconazole activates FgHog1, which in turn upregulates ergosterol biosynthetic genes through phosphorylation of FgSR (Liu et al., 2019). This study explored the transcriptional regulatory mechanism of *CsErg5B* and identified a novel cascade: HOG MAPK - CsAtf1 - *CsErg5B.* This fills a gap in *C. siamense* ergosterol gene regulation, extends the conserved role of the HOG MAPK pathway in ergosterol metabolism, and reveals species-specific regulatory features distinct from those in yeast and other fungi.

Our genetic complementation assays further defined the functional hierarchy of the HOG MAPK - CsAtf1 - *CsErg5B* cascade. Overexpression of *CsErg5B* partially restored conidial germination and fludioxonil sensitivity in both the Δ*CsAtf1* and Δ*CsPbs2* backgrounds, firmly establishing *CsErg5B* as a critical downstream effector of the HOG MAPK pathway. Notably, however, neither complementation strain recovered normal appressorium formation or full virulence (Fig S7). These observations indicate that this regulatory module mediates only a subset of biological processes rather than exerting global control. In line with our previous integrated RNA- Seq and ChIP- Seq analyses, CsAtf1 directly targets a large set of genes genome- wide, including 24 cytochrome oxidase family genes, among which *CsErg5B* is one direct target experimentally verified in this study (Guan et al., 2022). This explains why restoring *CsErg5B* expression only rescues defects in germination and fungicide sensitivity. We therefore infer that additional CsAtf1 target genes remain to be characterized, and these are likely responsible for regulating appressorium development, host infection, and pathogenicity.

Although we have clarified the role of the HOG MAPK - CsAtf1 - *CsErg5B* module in conidial germination, the underlying mechanisms remain unresolved. One key question is how CsAtf1 transcriptionally regulates *CsErg5B*, and the core binding motif of CsAtf1 on the *CsErg5B* promoter remains to be identified. Additionally, the molecular mechanism by which *CsErg5B* mediates conidial germination is still unclear. As a cytochrome oxidase-related gene involved in ergosterol biosynthesis, it remains elusive how *CsErg5B*-modulated ergosterol levels and membrane lipid composition control the initiation and progression of conidial germination. Notably, since *CsErg5B* overexpression restored both conidial germination and fludioxonil sensitivity, it is critical to examine whether these two processes are intrinsically linked. Our results support that the HOG MAPK - CsAtf1 - *CsErg5B* module is involved in regulating both traits, yet it remains unclear whether they are coordinately regulated through the same downstream pathway of *CsErg5B* or independently controlled by distinct mechanisms. These unanswered questions not only restrict our understanding of the HOG MAPK - CsAtf1 - *CsErg5B* regulatory network, but also highlight important directions for future research on the molecular basis of conidial germination and fungicide sensitivity in plant pathogenic fungi.

In summary, our findings reveal the HOG MAPK - CsAtf1 - *CsErg5B* cascade as a critical regulatory axis governing conidial germination and fludioxonil sensitivity in *C.siamense* (Fig 8),. CsAtf1 activates *CsErg5B* expression, which positively regulates conidial germination and enhances fludioxonil sensitivity, while repressing *CsCyp51G1* to suppress conidial development and reduce fungicide sensitivity. Genetic epistasis analysis revealed that *CsErg5B* is the dominant effector, with *CsCyp51G1* playing a secondary role. The working model links HOG MAPK signaling to ergosterol homeostasis providing a molecular framework for understanding infection-related morphogenesis and guiding anthracnose control strategies.

**Fig 8.**
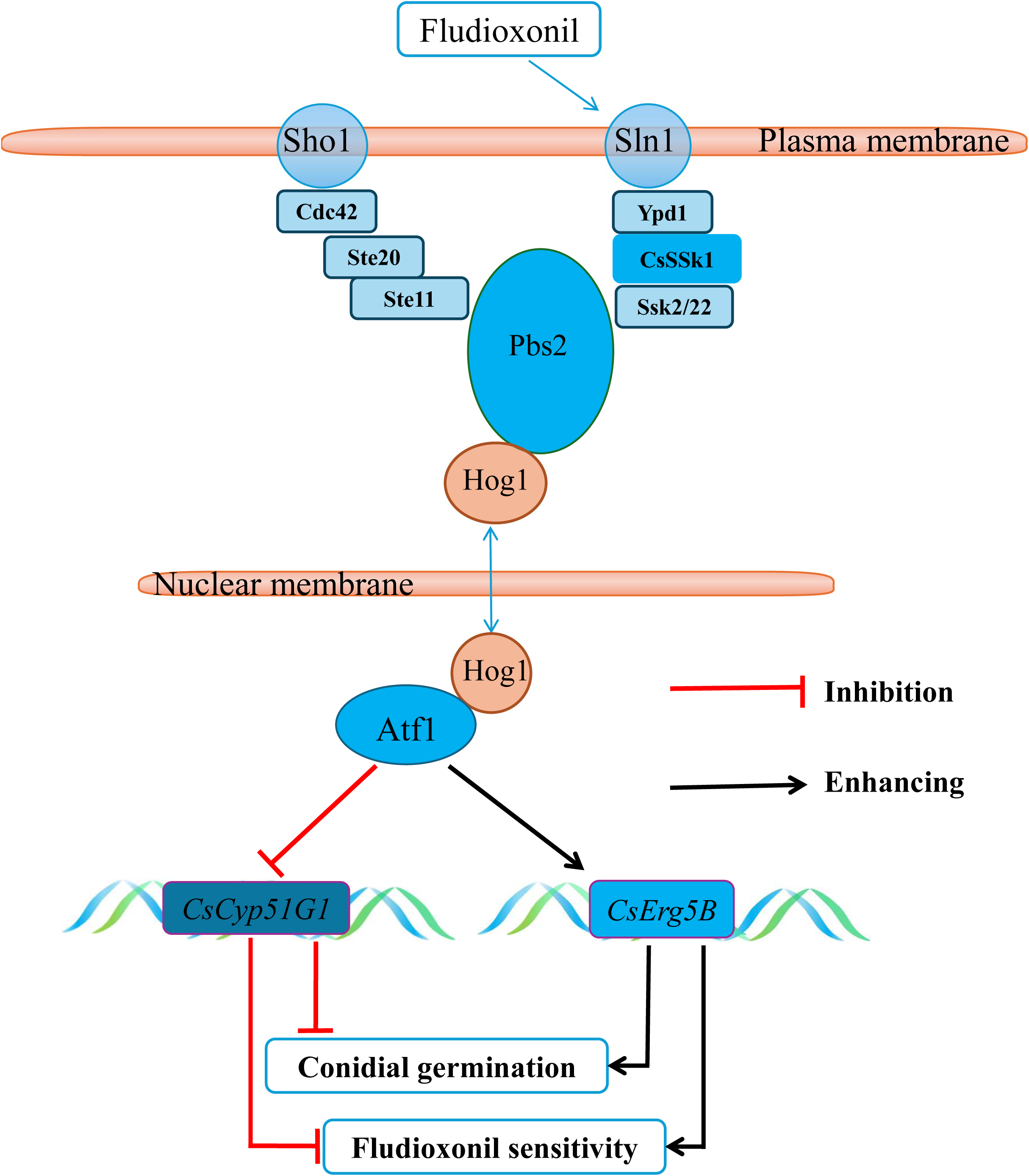
A proposed working model of the core HOG MAPK - CsAtf1 - *CsErg5B/CsCyp51G1* module regulating conidial germination and fludioxonil sensitivity. HOG MAPK- CsAtf1 activates *CsErg5B* and inhibits *CsCyp51G1* expression, thereby increasing conidial germination and fludioxonil sensitivity. *CsErg5B* serves as the primary downstream effector of CsAtf1 in controlling conidial development and fludioxonil sensitivity.

## 4 Materials and Methods

### 4.1 Strains, Culture Conditions and Chemicals

The *C. siamense* strain HN08, isolated from rubber tree, was used as the wild type (WT) in this study. Mutants Δ*CsPbs2*, Δ*CsAtf1* and other related transformants were derived from HN08 in this work or constructed earlier by our laboratory (Lin et al., 2018; Song et al., 2022). To collect conidia, mycelial blocks were incubated on PDA medium (Potato Dextrose Agar, Potato 200 g/L, Dextrose 20 g/L, and Agar 20 g/L) at room temperature under constant illumination for 3-5 days. For DNA, RNA, and total protein extraction, mycelia were harvested after incubation in liquid CM medium (0.6% yeast extract, 0.1% casein acid hydrolysate and 1% sucrose) at 28 °C in darkness for 2-3 days. Two DMI fungicides, difenoconazole (95% a.i.; Adama agricultural solutions Ltd.), tebuconazole (95% a.i.; Bayer CropScience, NC), and one phenylpyrrole fungicide, fludioxonil (95% a.i.; Bayer CropScience, NC) were used for sensitivity assays.

### 4.2 Verification of *CsErg5B* as a Direct Target of CsAtf1

#### 4.2.1 *CsErg5B* Gene Cloning and Sequence Analysis

The target gene sequence was identified by local BLAST analysis against the *C. siamense* HN08 whole-genome database, using a query sequence (Gene ID: 43608464). The primer pair *CsErg5B*-F/*CsErg5B*-R was designed for this study and is provided in Table S1. Both genomic DNA and cDNA fragments of *CsErg5B* were amplified and subjected to Sanger sequencing. Conserved domains were predicted with the Simple Modular Architecture Research Tool (SMART; http://smart.embl-heidelberg.de, accessed 2 April 2024). Multiple sequence alignment was carried out using the ClustalW program (Larkin et al., 2007). Phylogenetic analysis was performed using the maximum-likelihood algorithm implemented in MEGA6 software (Tamura et al., 2013), with branch confidence estimated by 1000 bootstrap replications.

#### 4.2.2 Yeast One-Hybrid (Y1H) Assays

Yeast one- hybrid (Y1H) assays were performed to verify the interaction between CsAtf1 and the *CsErg5B* promoter. The promoter fragment of *CsErg5B* was ligated into the *Eco*RI and *Sac*I restriction sites of the pHIS2 backbone, yielding the reporter construct pHIS2-P- *CsErg5B*. Meanwhile, the full-length coding region of *CsAtf1* was inserted into the *Bam*HI and *Eco*RI sites of pGADT7 to generate the effector plasmid pGADT7- *CsAtf1*. The recombinant plasmids were co-transformed into yeast strain Y187. Positive transformants were cultivated on SD/-Trp/-Leu medium and SD/-Trp/-Leu/-His medium containing 80 mM 3-AT. The pGADT7- 53/pHIS2- 53 combination was employed as a positive control, whereas co-transformation of the empty pGADT7 and pHIS2 vectors served as a negative control. All culture plates were maintained at 28 °C for 3 to 4 days.

#### 4.2.3 Electrophoretic Mobility Shift Assay (EMSA)

Recombinant CsAtf1 protein was expressed in a prokaryotic system. The coding sequence of *CsAtf1* was inserted into the pET-32a vector with an N-terminal 6×His tag to construct the expression plasmid pET-32a-*CsAtf1*. The His-tagged CsAtf1 fusion protein was induced and expressed in *Escherichia coli* BL21 cells, followed by affinity purification. Target DNA fragments were amplified by PCR, purified using a Gel Extraction Kit (Tiangen, P.R. China), and labeled with biotin via an EMSA Probe Biotin Labeling Kit (Beyotime, P.R. China) according to the manufacturer’s instructions. The binding reaction was performed using a Chemiluminescent EMSA Kit (Beyotime, P.R. China). Protein–DNA complexes were separated on a 4% non-denaturing polyacrylamide gel in 0.5×TBE buffer, transferred to a nylon membrane, and cross-linked by UV irradiation. Finally, chemiluminescent signals were detected in accordance with the kit manual. Additional experimental procedures were performed as previously described (Guan et al., 2022).

#### 4.2.4 Total RNA Extraction and Quantitative Real-time pCR (qRT-PCR) Analysis of *CsErg5B* Expression

Mycelia were collected from wild-type HN08, the Δ*CsAtf1* mutant, and the Δ*CsAtf1*/*CsAtf1* complemented strain for total RNA extraction using the RNAprep Pure Plant Kit (Tiangen, Beijing, China). First-strand cDNA was generated from 1 μg of purified RNA with the TransScript One-Step gDNA Removal and cDNA Synthesis SuperMix (TransGen Biotech, Beijing, China). Transcriptional levels of *CsErg5B* were examined by quantitative real-time PCR (qRT-PCR) with an ABI 7500 Real-Time PCR System (Applied Biosystems, Waltham, MA, USA). Each amplification reaction was performed in a 10 μL volume containing SYBR Premix Dimer Eraser (Takara, Beijing, China). The transcriptional expression levels of *CsAtf1* and *CsErg5B* were also detected in liquid cultures with or without treatment of anisomycin (2 × 10⁻³ μg/mL), SB203580 inhibitor (20 μg/mL), and fludioxonil (at a final concentration of 50 μg/mL). The *Colletotrichum* actin gene was used as an internal reference for normalization, and relative gene expression levels were calculated using the 2^⁻ΔΔCT^ method (Maren et al., 2023). Three independent biological replicates and at least three technical replicates were included for each sample. Primer sequences are shown in Table S1.

### 4.3 *CsErg5B* Gene Deletion and Verification

Targeted gene deletion was performed via homologous recombination as described previously (Guan et al., 2022). The upstream and downstream flanking regions of *CsErg5B* were amplified with primer pairs UF/UR and DF/DR, respectively, with UR and DF containing a linker for the *ILV1* chlorimuron resistance cassette. The two fragments were fused with the *ILV1* cassette amplified from pCX62-S, and the full fusion construct was amplified with UF/DR and transformed into HN08 protoplasts as described previously (Song et al., 2022).

Putative transformants were verified by PCR, sequencing and Southern blotting. Four primer pairs were applied: *CsErg5B*-F/R for full-length gene amplification, *CsErg5B*-UF/*ILV*-R and *CsErg5B*-DR/*ILV*-F for junction fragment detection, and *CsErg5B*-OUF/*CsErg5B*-ODR for amplification of the flanking region. Sequencing confirmed that the endogenous *CsErg5B* was replaced by the *ILV1* marker. For Southern blotting, genomic DNA was isolated via the CTAB method (Sambrook et al., 1982), completely digested with *Eco*RI, and hybridized with a DIG-labeled probe specific to *ILV1*. Hybrid signals were detected using a DIG Luminescent Detection Kit (Roche, Mannheim, Germany) to confirm targeted gene deletion.

### 4.4 *CsErg5B* Gene Overexpression

#### 4.4.1 Construction of Δ*CsErg5B*/*CsErg5B*-OE Strain

To construct the Δ*CsErg5B*/*CsErg5B*-OE overexpression strain, the pXY203 vector harboring the *HPH* hygromycin resistance cassette and *RP27* promoter was linearized by restriction digestion. The full-length *CsErg5B* coding sequence was amplified by PCR. The linearized vector and target fragment were co-transformed into yeast strain XK1-25, and positive clones were selected on SD/-Trp medium. The recombinant plasmid pXY203-*RP27*-*CsErg5B* was isolated from yeast, amplified in *Escherichia coli*, and then transformed into HN08 protoplasts. Putative transformants were selected on PDA medium supplemented with 1 M sucrose (PDS) containing 600 µg/mL hygromycin. The validated strain was designated.

#### 4.4.2 Construction of Δ*CsAtf1*/*CsErg5B*-OE Strain

To generate the Δ*CsAtf1*/*CsErg5B*-OE overexpression strain, the *CsErg5B* cDNA was amplified with primers pBAR-*CsErg5B*-F/pBAR-*CsErg5B*-R and cloned into the pBAR-GFP vector carrying the *gpdA* promoter and *HPH* gene. The recombinant plasmid pBAR-*CsErg5B*-GFP was transformed into protoplasts of the Δ*CsAtf1* mutant (chlorimuron ethyl-resistant knockout strain) to obtain the Δ*CsAtf1*/*CsErg5B*-OE strain (Song et al., 2022).

#### 4.4.3 Construction of Δ*CsPbs2*/*CsErg5B*-OE Strain

To generate the Δ*CsPbs2*/*CsErg5B*-OE overexpression strain, the *CsErg5B* cDNA was amplified with primers pKNT-*CsErg5B*-F/pKNT-*CsErg5B*-R and cloned into the pKNT-GFP vector harboring the *RP27* promoter and *G418* resistance gene. The recombinant plasmid pKNT-*CsErg5B*-GFP was transformed into Δ*CsPbs2* mutant protoplasts (hygromycin-resistant knockout strain), and positive transformants were screened on TB3 medium (3 g/L yeast extract, 3 g/L casamino acids, 200 g/L sucrose, 20 g/L agar) supplemented with 800 μg/mL G418, yielding the Δ*CsPbs2*/*CsErg5B-OE* overexpression strain (Guan et al., 2022).

#### 4.4.4 Validation of *CsErg5B* Overexpression Strains

The relative expression levels of the *CsErg5B* gene in the gene deletion mutants (Δ*CsAtf1* and Δ*CsPbs2*) and overexpression strains (Δ*CsErg5B*/*CsErg5B*-OE, Δ*CsAtf1*/*CsErg5B*-OE, and Δ*CsPbs2*/*CsErg5B*-OE) were determined by quantitative real-time PCR (qRT-PCR). All reactions were performed in technical triplicate for three independent biological replicates. Primer sequences used in this study are provided in Table S1.

### 4.5 Construction of Δ*CsErg5B*/Δ*CsCyp51G1* Double Deletion Mutant

To generate the Δ*CsErg5B*/Δ*CsCyp51G1* double deletion mutant, the *CsCyp51G1* deletion cassette was introduced into the Δ*CsErg5B* mutant background. The upstream and downstream flanking regions of the *CsCyp51G1* gene were amplified using the primer pairs *CsCyp51G1*-UF/*CsCyp51G1*-UR and *CsCyp51G1*-DF/*CsCyp51G1*-DR, respectively; *CsCyp51G1*-UR and *CsCyp51G1*-DF contained linker sequences for the hygromycin resistance gene *HPH*. Using the Split-Marker method, the two flanking fragments were separately fused with the split *HPH* gene fragments (amplified from plasmids PCSN43 and PBAR-GFP ) to obtain the left and right fragments (Garcia et al., 2023). Subsequently, the two fusion fragments were introduced into protoplasts of the Δ*CsErg5B* mutant and screened on PDS medium supplemented with hygromycin. Putative double deletion mutants were verified by PCR using primer pairs specific to both gene loci. The confirmed strain was designated Δ*CsErg5B*/Δ*CsCyp51G1* and used for subsequent phenotypic assays.

### 4.6 Phenotype analysis

#### 4.6.1 Hyphal Growth, Conidial Morphology, Sporulation, Conidial Germination and Appressorium Formation

Conidial suspensions of HN08 wild-type, Δ*CsErg5B* and Δ*CsErg5B*/*CsErg5B*-OE strains were adjusted to 1 × 10⁵ conidia/mL. For growth assays, 10 μL aliquots were inoculated on CM plates and cultured at 28 °C for 5 days before colony diameter measurement. For morphology observation, 20 μL suspensions were spotted onto glass slides and examined microscopically. For sporulation assessment, 10 μL suspensions (1 × 10⁵ conidia/mL) of the above strains plus Δ*CsAtf1* and Δ*CsAtf1*/*CsErg5B*-OE were inoculated on PDA plates. After 5 days at 28 °C, conidia were harvested with sterile ddH₂O, filtered, diluted to 1 mL, and counted with a hemocytometer. For germination and appressorium formation assays, 20 μL suspensions of all tested strains (including Δ*CsAtf1*/*CsErg5B*-OE and Δ*CsPbs2*/*CsErg5B*-OE) were inoculated. Germination was recorded at 0, 2, 4, 6, and 8 h, and appressorium formation at 16 h. At least 300 conidia were counted per assay with three biological replicates.

#### 4.6.2 Pathogenicity Assays

The virulence of the wild-type (WT) strain, Δ*CsAtf1* mutant, Δ*CsErg5B* mutant, Δ*CsErg5B*/*CsErg5B*-OE strain and Δ*CsAtf1*/*CsErg5B*-OE strain was evaluated on detached light-green rubber tree (CATAS 7-33-97) leaves in vitro. Conidial suspensions (adjusted to 1 × 10^6^ conidia/mL, 10 μL per drop) were spotted onto detached young leaves that were either wounded with a sterile needle or left unwounded. Four drops were inoculated on both sides of the main vein per leaf, corresponding to the WT strain, deletion mutant, overexpression transformant and sterile ddH₂O (control), respectively. Each strain was prepared with at least 30 biological replicates. The inoculated leaves were incubated at 28 ℃ in darkness for 24 h, then transferred to light conditions. Disease development was observed and lesion areas measured at 3, 4, and 5 days post inoculation.

#### 4.6.3 Osmotic Stress and Fungicide Sensitivity Assays

Conidial suspensions (1 × 10⁶ conidia/mL) of *C. siamense* WT, Δ*CsErg5B* and Δ*CsErg5B*/*CsErg5B*-OE strains were inoculated onto CM plates supplemented with 0.6 M NaCl, 0.7 M KCl, 50 μg/mL Congo Red, 1 M glucose, 1 M sorbitol or 2.5 mM H₂O₂, with untreated plates as control. A 10 μL suspension was spotted per plate. All assays were performed in triplicate across three independent experiments. Following 7 days of incubation at 28 ℃, colony diameters were measured and photographed on days 3, 5 and 7.

For fungicide sensitivity assays, the same conidial preparation was used. Sensitivity to tebuconazole (0, 0.25, 0.5, 1, 2 μg/mL) and difenoconazole (0, 0.01, 0.1, 0.5, 1 μg/mL) was determined in WT, Δ*CsErg5B* and Δ*CsErg5B*/*CsErg5B*-OE, while sensitivity to fludioxonil (0, 0.01, 0.1, 0.5, 5, 10 μg/mL) was tested in all strains (WT, Δ*CsErg5B*, Δ*CsErg5B*/*CsErg5B*-OE, Δ*CsAtf1*, Δ*CsAtf1*/*CsErg5B*-OE, Δ*CsPbs2*, Δ*CsPbs2*/*CsErg5B*-OE). A 10 μL aliquot was inoculated onto CM plates with or without fungicides (3 replicates per concentration). Colony diameters were measured after 3-7 days at 28 °C in darkness, and growth inhibition rates were calculated.

#### 4.6.4 Ergosterol Content Determination and Exogenous Ergosterol Supplementation Assay

The ergosterol content of WT HN08, Δ*CsAtf1*, and Δ*CsErg5B* strains was determined by a commercial service using high-performance liquid chromatography (HPLC). HPLC analysis was performed using a Waters 2695 system with a Compass C18(2) reversed-phase column (250 mm × 4.6 mm, 5 μm) at 25 °C with a flow rate of 1 mL/min. The mobile phase was methanol (100%). The detection wavelength was 282 nm, injection volume was 10 µL, and retention time was approximately 12 min. A standard curve was established using ergosterol standard (purity ≥ 98%, Yuanye) at concentrations ranging from 0.5 to 80 ppm. Data are means of three replicates.

Exogenous ergosterol was added to conidial suspensions at a final concentration of 12.5 μg/mL. Conidial germination was observed and recorded at 0, 2, 6, and 8 h. Conidial suspensions without ergosterol supplementation were used as controls. All experiments were performed in three biological replicates.

## Supporting Informaion

Additional supporting information can be found online in the Supporting Information section.

**Fig S1.**
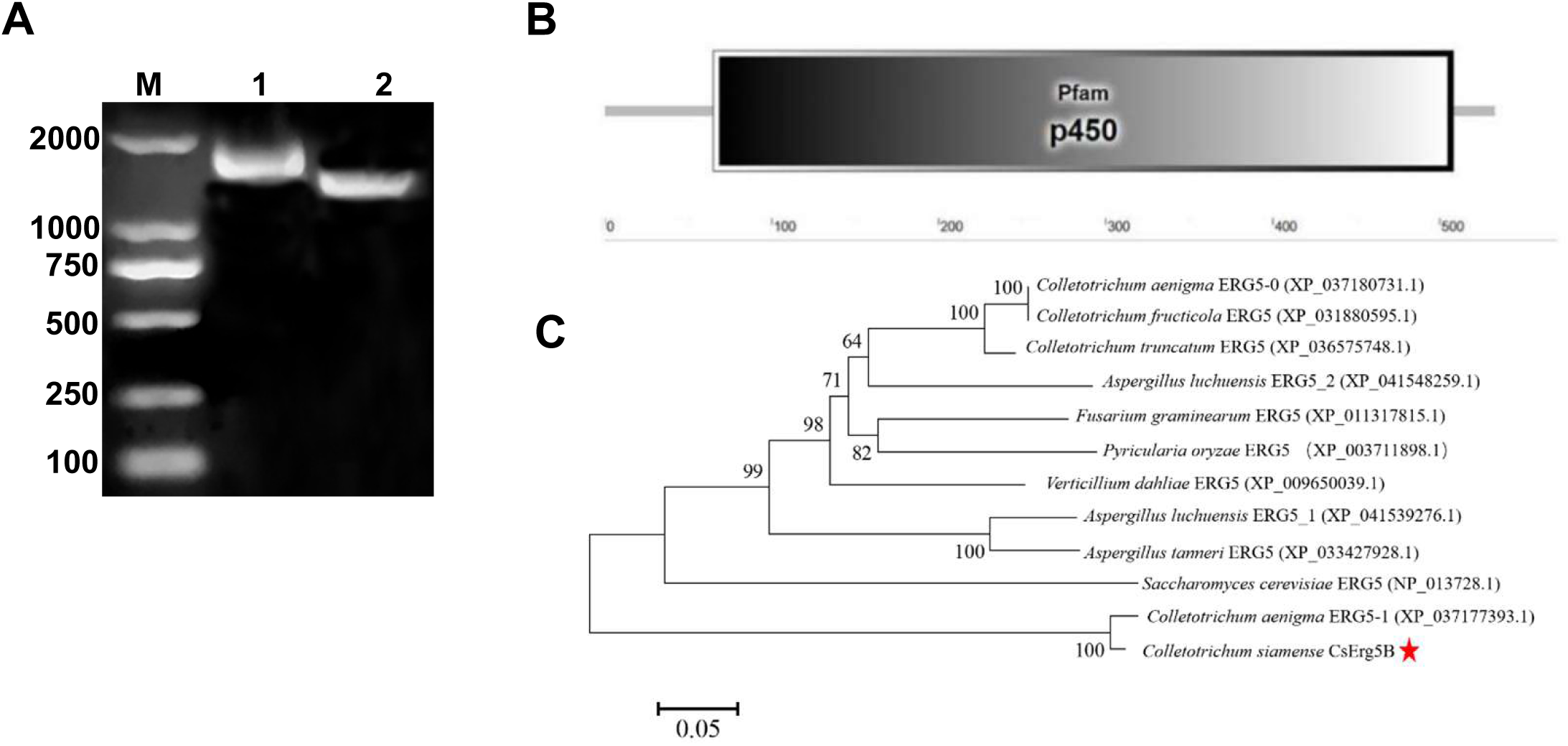
Cloning and sequence analysis of the *CsErg5B* gene. **(A)** PCR amplification of *CsErg5B* DNA (lane 1) and cDNA (lane 2) fragments using primer pair *CsErg5B*-F/*CsErg5B*-R, resolved by agarose gel electrophoresis. **(B)** Protein domain architecture of CsErg5B in *C. siamense*. **(C)** Phylogenetic analysis of CsErg5B and its homologous proteins based on amino acid sequences.

**Fig S2.**
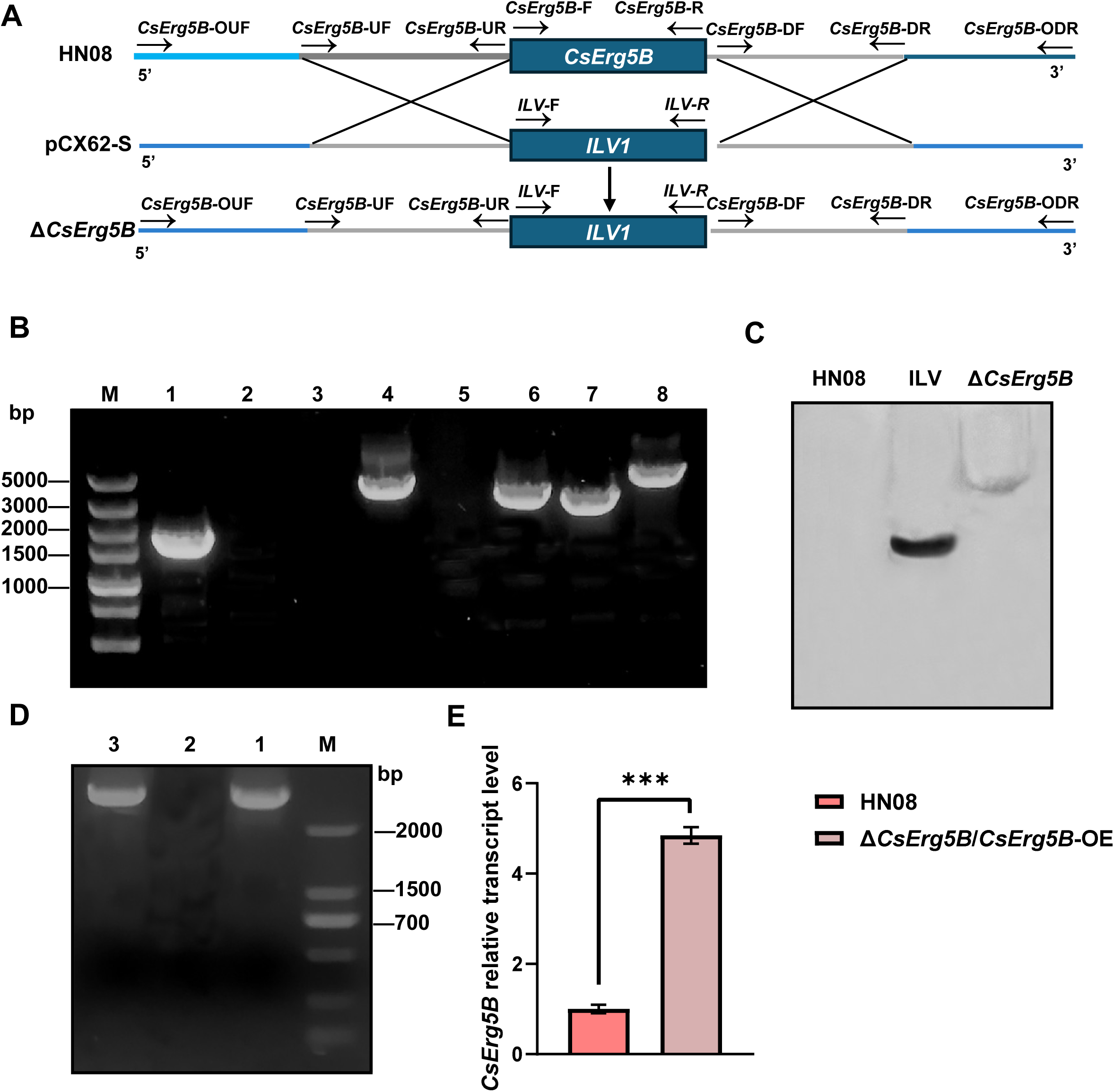
Construction and molecular validation of the Δ*CsErg5B* knockout mutant and Δ*CsErg5B*/*CsErg5B*-OE overexpression strain. **(A)** Schematic diagram of Δ*CsErg5B* knockout construction. **(B)** PCR verification of the Δ*CsErg5B* mutant. Lane M indicates the DL5000 DNA marker. Lanes 1-4 were amplified from genomic DNA of the wild type strain HN08, and lanes 5-8 were amplified from Δ*CsErg5B* mutant genomic DNA. The primer pairs *CsErg5B*-F/R, *CsErg5B*-UF/*ILV1*-R, *ILV1*-F/*CsErg5B*-DR, and *CsErg5B*-OUF/*CsErg5B*-ODR were used for detection in lanes 1/5, 2/6, 3/7, and 4/8, respectively. **(C)** Southern blot analysis of the Δ*CsErg5B* mutant. Genomic DNA was digested with *Eco*RI, and the *ILV1* gene fragment was used as a specific probe. A single hybridizing band was present in the Δ*CsErg5B* mutant, while no signal was detected in the wild-type strain HN08. **(D)** PCR verification of the Δ*CsErg5B*/*CsErg5B*-OE strain. Lane M, DNA DL2000 marker. The primer pair *RP27*-F/*CsErg5B*-R was used for amplification in lanes 1–3. The PCR templates of lane 1, lane 2 and lane 3 were the overexpression vector pXY203-*RP27*-*CsErg5B*, genomic DNA of wild-type strain HN08, and genomic DNA of the Δ*CsErg5B*/*CsErg5B*-OE strain, respectively. **(E)** Transcription level analysis of *CsErg5B* in the Δ*CsErg5B*/*CsErg5B*-OE strain. Data are presented as mean ± SD. *** represents an extremely significant difference (*p* < 0.001).

**Fig S3.**
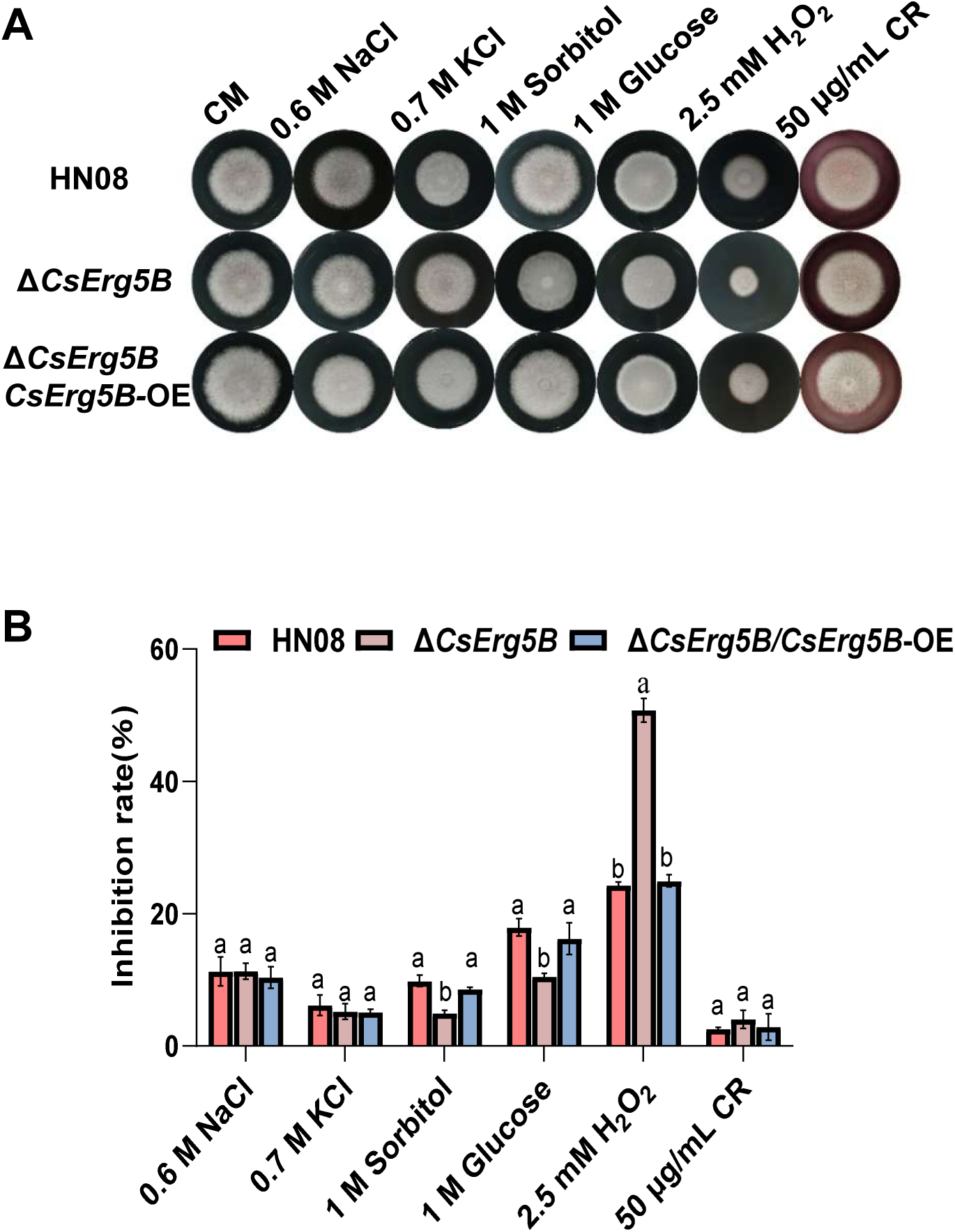
The effects of *CsErg5B* deletion on stress responses.. (**A, B)** Mycelial growth of the wild-type HN08, Δ*CsErg5B*, and Δ*CsErg5B*/*CsErg5B*-OE under various stress conditions, including salt stress (NaCl/KCl), cell wall stress (Congo Red), osmotic stress (sorbitol/glucose), and oxidative stress (H₂O₂).

**Fig S4.**
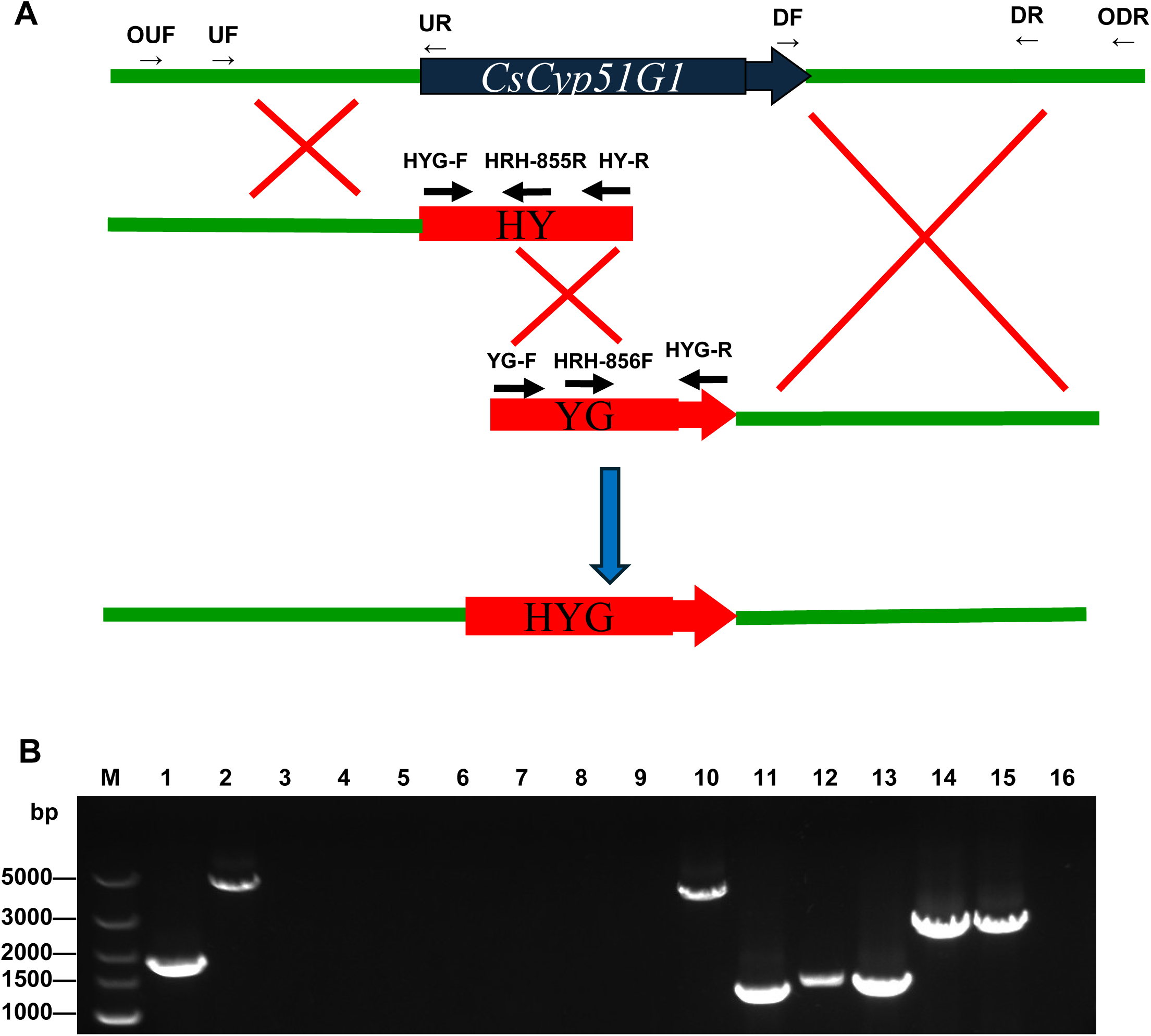
Construction and molecular validation of double-gene deletion mutant Δ*CsErg5B*/Δ*CsCyp51G1*. **(A)** Schematic diagram of double-gene knockout. **(B)** PCR identification of the Δ*CsErg5B* single mutant and Δ*CsErg5B*/Δ*CsCyp51G1* double mutant. Genomic DNA of the Δ*CsErg5B* strain was used as PCR templates for lanes 1-8, and the Δ*CsErg5B/*Δ*CsCyp51G1* strain templates were applied for lanes 9-16. Multiple primer pairs were used for mutant verification: *CsCyp51G1*-F/R (lanes 1, 9), *CsCyp51G1*-OUF/ODR (lanes 2, 10), *CsCyp51G1*-OUF/*HRH*-855R (lanes 3, 11), *HRH*-856F/*CsCyp51G1*-ODR (lanes 4, 12), *HYG*-F/R (lanes 5, 13), *CsCyp51G1*-OUF/ *HYG*-R (lanes 6, 14), *HYG*-F/*CsCyp51G1*-ODR (lanes 7, 15), and *CsErg5B*-F/R (lanes 8, 16). Target fragments with expected sizes and negative amplification results confirmed the successful construction of the double-gene deletion mutant.

**Fig S5.**
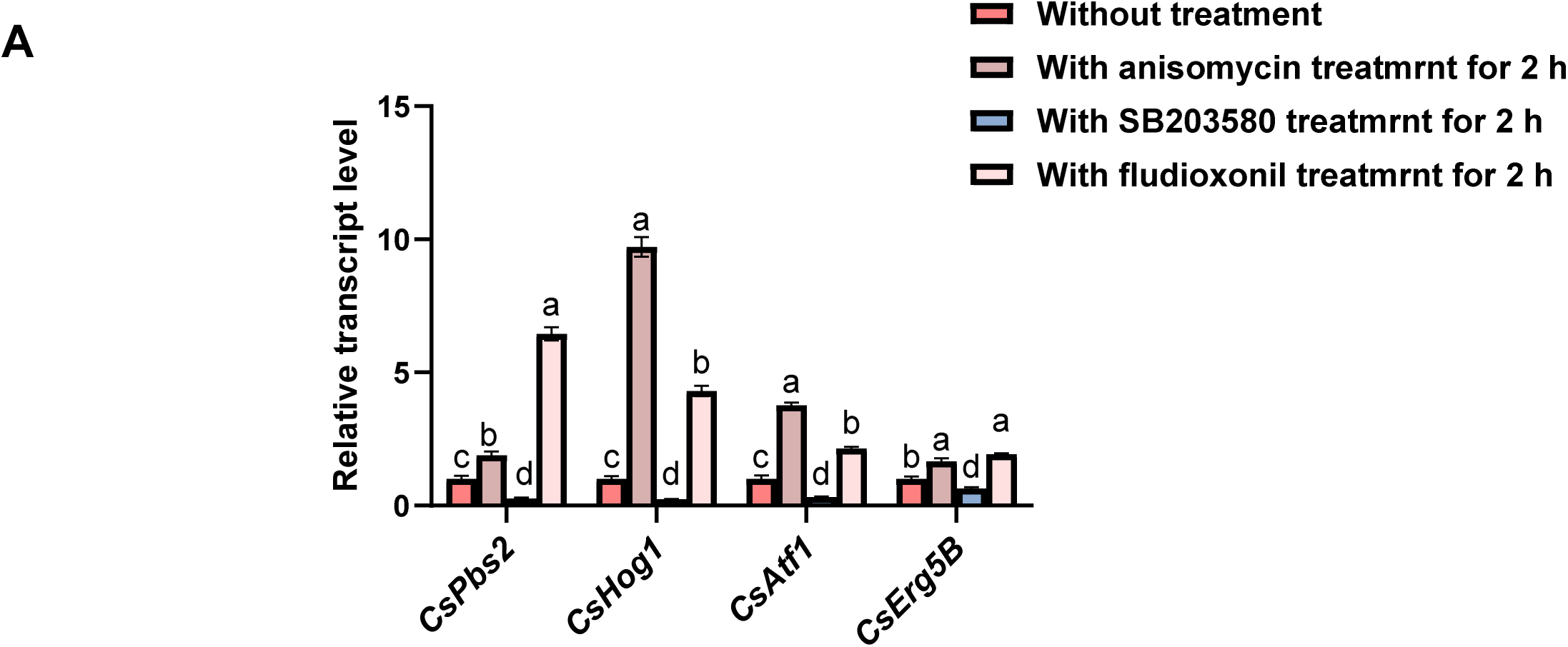
Expression levels of *CsPbs2*, *CsHog1*, *CsAtf1* and *CsErg5*B in HN08 with or without treatments of 2×10^−3^ μg/mL anisomycin (a HOG MAPK activator), 20 μg/mL SB203580 (a p38 MAPK inhibitor), and 50 μg/mL fludioxonil (a phenylpyrrole fungicide). Mycelia of the wild-type strain HN08 were treated with the HOG MAPK activator anisomycin, the p38 MAPK inhibitor SB203580, and fludioxonil, respectively. The transcript levels of *CsPbs2*, *CsHog1*, *CsAtf1* and *CsErg5B* were detected by qRT- PCR after 2 hpi of different treatments. Data are mean ± SD. Different letters indicate significant differences at *p* < 0.05.

**Fig S6.**
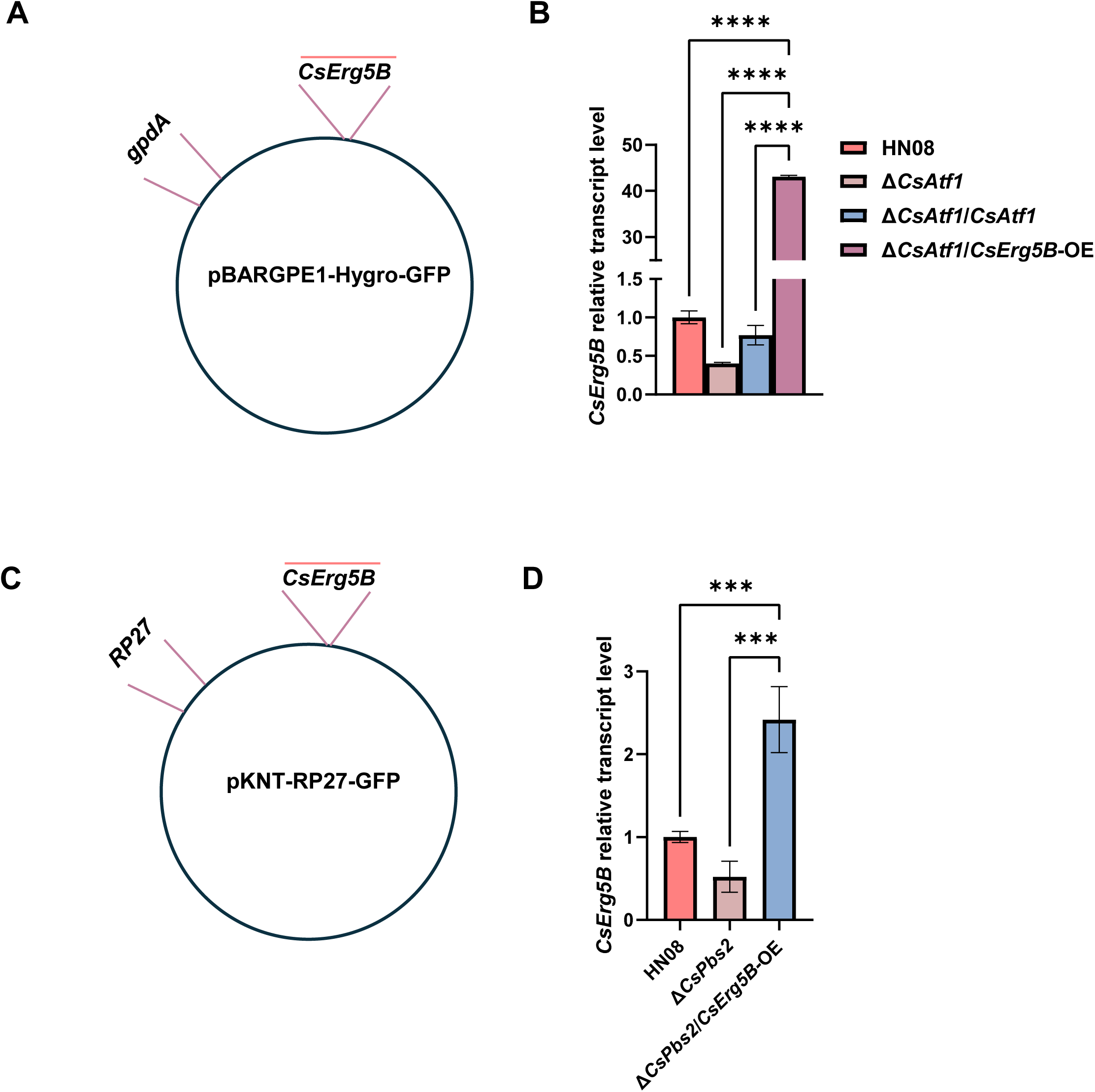
Construction of the Δ*CsAtf1*/*CsErg5B*-OE and Δ*CsPbs2*/*CsErg5B*-OE strains. **(A, C)** Schematic diagrams of vector construction. **(B, D)** The relative expression levels of target genes in the wild type, deletion mutants and overexpression strains determined by qRT-PCR. Data are presented as mean ± SD. *** represents an extremely significant difference (*p* < 0.001), and **** represents an ultra-significant difference (*p* < 0.0001).

**Fig S7.**
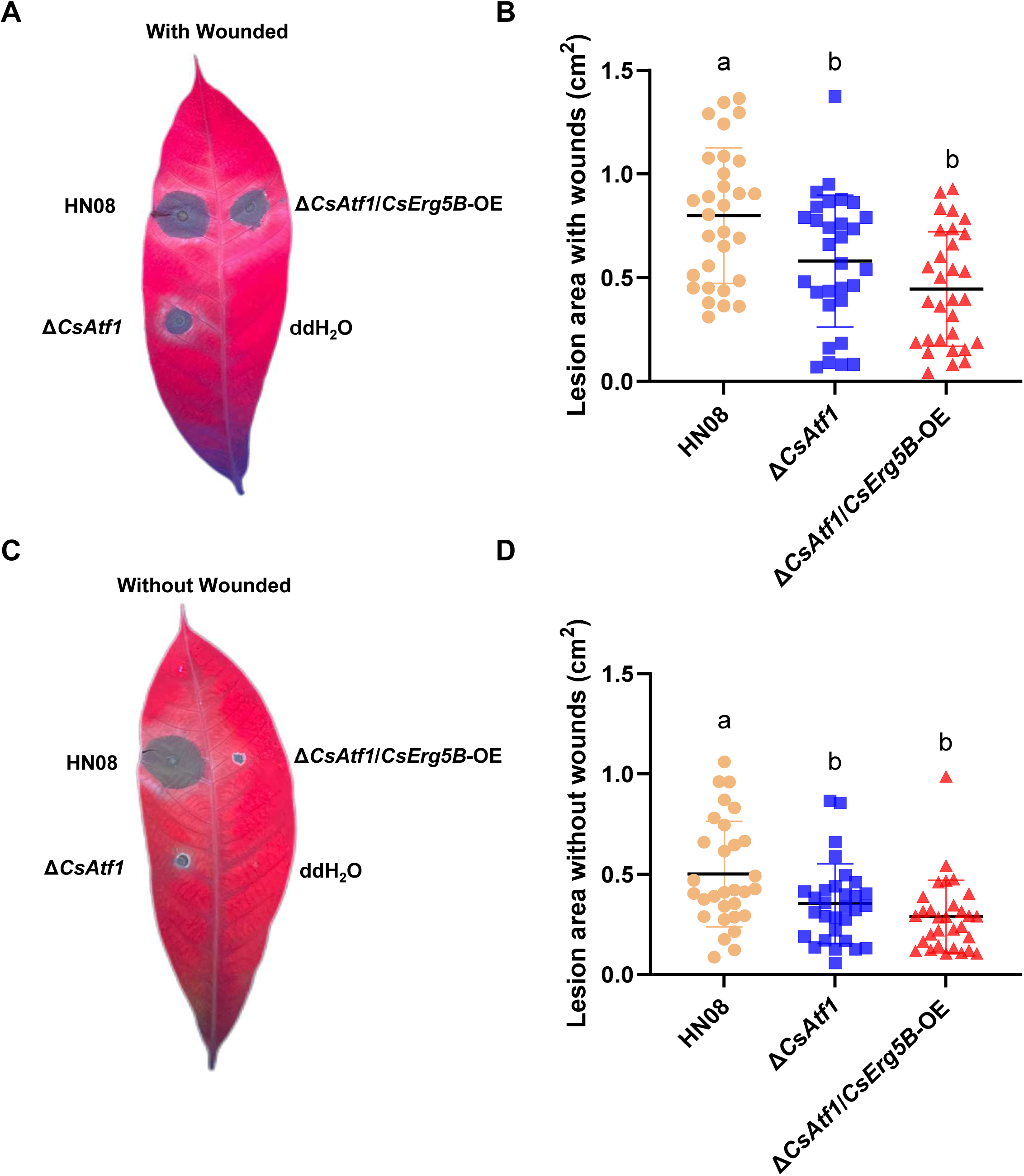
Pathogenicity assay of the wild-type strain HN08, Δ*CsAtf1* mutant, and Δ*CsAtf1*/*CsErg5B*-OE strain. **(A, B)** Disease symptoms and statistical analysis of lesion areas on wounded leaves at five days post-inoculation. **(C, D)** Disease symptoms and statistical analysis of lesion areas on unwounded leaves at five days post-inoculation. Overexpression of *CsErg5B* could not rescue the reduced pathogenicity of the Δ*CsAtf1* mutant, with obvious virulence differences between the wild type and related mutants. Data are presented as mean ± SD, and different letters represent significant difference at *p* < 0.05.

**Table S1.**
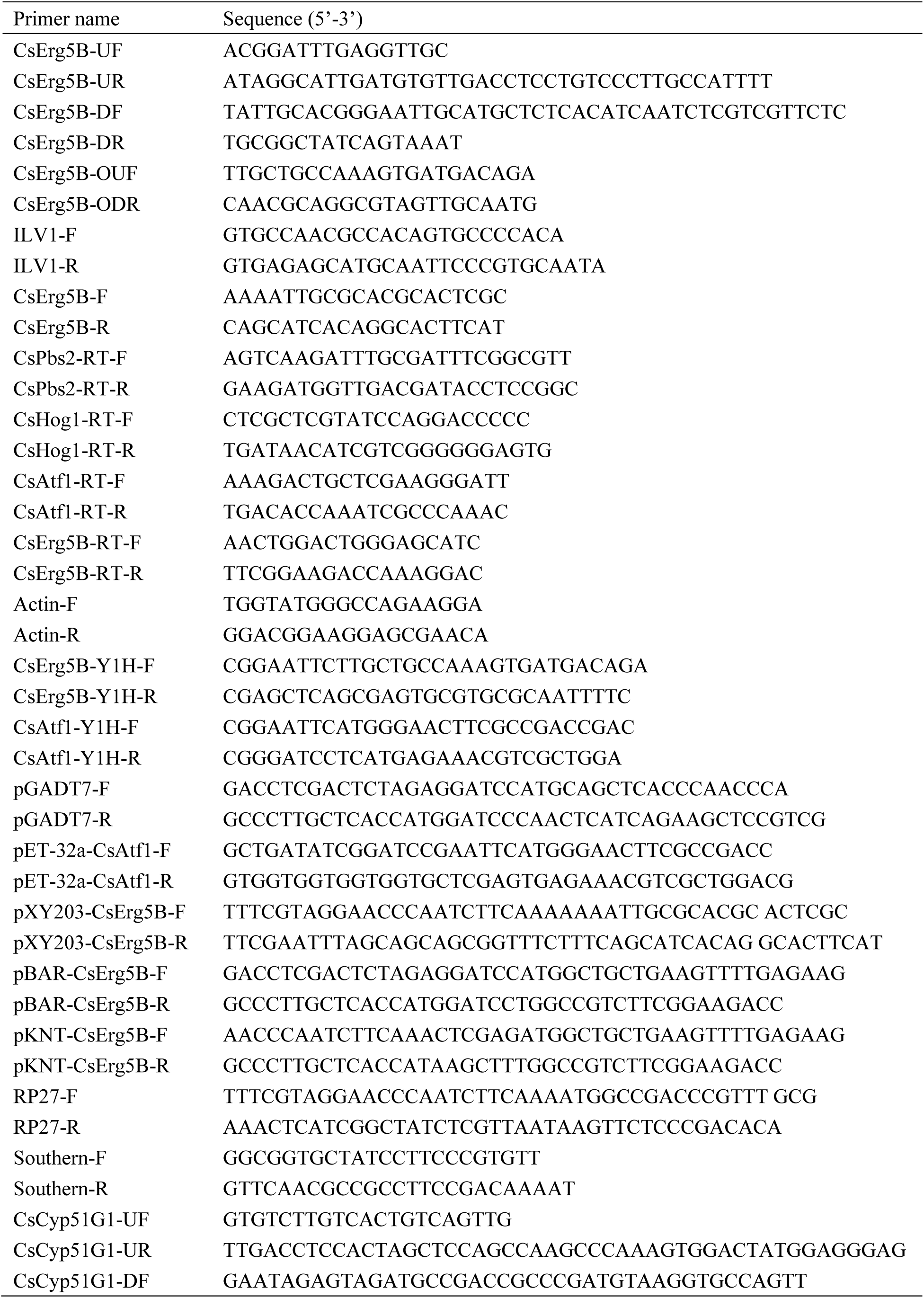
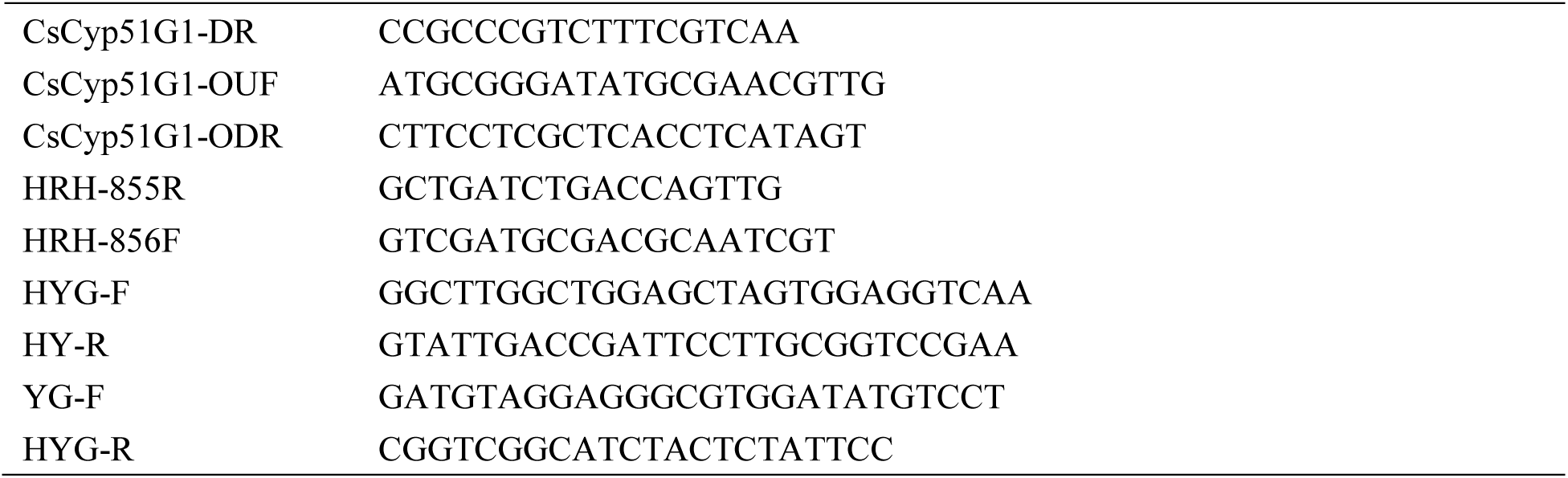
List of primers used in the study.

## Author contributions

**Yuqing Lin**: Writing - original draft, Investigation, Data curation. **Kuaikuai Wang**: Validation, Investigation. **Xiaoling Guan**: Investigation; **Miao Song**: Methodology. **Zhenyu Han**: Formal analysis. **Wenbo Liu**: Data curation. **Wei Wu**: Writing—review and editing. **Yu Zhang:** Funding acquisition. **Weiguo Miao**:Funding acquisition. **Chunhua Lin**: Supervision, Writing—review and editing, Project administration, Funding acquisition.

## DATA AVAILABILITY

The data that support the findings of this study are available within the manuscript and its Supporting Information files. Additional information is available from the corresponding authors upon reasonable request.

